# V-ATPase-Driven Lysosomal Activation Orchestrates MEK2-Induced Endothelial Reprogramming

**DOI:** 10.64898/2026.06.09.731136

**Authors:** Ziad Sabry, Max Keller, Liu Liu, Morgan Salmon, Zhong Wang

## Abstract

Direct lineage reprogramming holds therapeutic promise but often depends on transcription factor overexpression, resulting in unstable phenotypes. Here, we describe a novel strategy to convert fibroblasts into endothelial-like cells by activating lysosomal activity. Constitutively active MEK2 induces an endothelial gene program via sustained MAPK/ERK signaling, leading to enhanced vacuolar ATPase (V-ATPase) activity, lysosomal acidification, extracellular matrix degradation, and angiogenic behavior. V-ATPase inhibition impairs these effects, whereas pharmacologic activation with EN6 recapitulates key features of reprogramming and promotes nuclear translocation of TFEB, a master lysosomal regulator. Consistently, TFEB overexpression–particularly a phospho-deficient mutant–boosts lysosomal function and endothelial gene expression. These findings define a MAPK–V-ATPase–TFEB axis that drives endothelial reprogramming and highlight the lysosome as a central hub for cell fate transitions, offering an organelle-centric framework for regenerative medicine.

**Highlights:**

- Sustained MEK2 activation reprograms fibroblasts into endothelial-like cells
- MEK2 enhances lysosomal acidification by upregulating V-ATPase subunits
- V-ATPase drives acidification and ECM remodeling for endothelial reprogramming
- V-ATPase activation promotes TFEB nuclear entry and endothelial gene expression

## Introduction

Cardiovascular disease is the leading cause of mortality worldwide. In ischemic heart disease, such as myocardial infarction (MI), obstruction of coronary blood flow leads to cardiomyocyte death and fibrotic scar formation in the infarcted myocardium. The injured endothelium triggers inflammatory and fibrogenic responses, wherein surviving endothelial cells (ECs) can undergo endothelial-to-mesenchymal transition to become fibroblast-like cells, and paracrine signals activate resident fibroblasts into myofibroblasts.^1–3^ The resulting myofibroblasts deposit excess extracellular matrix (ECM), stiffening the cardiac tissue and impairing contractile function.^4–6^ Unlike regenerative organs, the adult heart has minimal endogenous stem cell activity to replenish the lost myocardium.

One promising approach to address the lack of endogenous stem cell therapy in the adult heart is direct lineage reprogramming—converting resident somatic cells (such as fibroblasts) into another cell type (such as endothelial cells) without passing through a pluripotent state.^7^ Several groups have demonstrated the conversion of fibroblasts into induced endothelial cells (iECs) using defined transcription factors; however, these iECs often exhibit immature or transient phenotypes.^8–13^ Increasing evidence suggests that successful cell fate change requires not only transcriptional rewiring but also coordination between cellular information processing and structural organization. Post-translational modifications (PTMs), such as phosphorylation, encode dynamic signaling information that shapes gene regulatory networks, while organelles and sub-organelle architectures provide the physical and metabolic context in which these signals are interpreted.^14–16^ Emerging work across developmental and reprogramming systems indicates that alterations in cell structure—spanning organelle function, intracellular trafficking, and local microenvironments—can actively influence lineage stability and plasticity, rather than passively responding to transcriptional changes.^17^ However, studies systematically linking PTMs and cell structure regulation to direct lineage reprogramming remain limited, leaving most mechanistic connections unexplored. Thus, cell fate transitions may be governed by an integrated information–structure framework, in which signaling states conveyed by PTMs interface with cell structure organization to stabilize or redirect cell identity. Leveraging this coordination may offer a means to overcome the incomplete or unstable phenotypes that have limited current transcription factor–based reprogramming strategies.

In this study, we aim to determine how PTMs and cell structure can coordinate to drive fibroblast conversion into iECs. We demonstrate that activating lysosomal function promotes EC-like reprogramming of MEFs. Our initial screen identifies sustained MAPK/ERK activation as a key driver of endothelial induction. Under this condition, we observe pronounced upregulation of V-ATPase, a central driver of lysosomal activity, which in turn mediates downregulation of fibroblast-associated ECM genes and promotes local ECM remodeling. Furthermore, direct activation of V-ATPase with the small molecule EN6 also enhances endothelial reprogramming, accompanied by nuclear translocation of the lysosomal transcription factor TFEB. These findings define a MAPK–V-ATPase–TFEB axis that drives endothelial reprogramming and position the lysosome as a master regulator—rather than a bystander—of cell fate change and a targetable organelle for regenerative intervention. This research represents one of the first demonstrations that organelle function can be directly harnessed to reprogram somatic cell identity. Our work suggests that organelle-based modulation may serve as a generalizable paradigm for cell fate engineering and regenerative therapy.

## Materials and Methods

### Assembly of endothelial drivers and kinases plasmids

To generate lentiviral particles, we utilized the EF1a-mCherry-P2A-hygro backbone, a plasmid gifted by Prashant Mali (Addgene plasmid #135003; http://n2t.net/addgene:135003; RRID). Genes of interest were amplified from their respective plasmids using the primers listed in **Table 1** and Taq DNA polymerase (Cat. No. M0267L, New England Biolabs, Ipswich, MA, USA). Gibson assembly was performed following the protocol described by Parekh et al. 2018.^18^ The lentiviral vectors used for overexpression of TFEB and its phosphorylation-resistant mutant TFEB^S141A/S210A^ (EF1A-TFEB-P2A-hygro and EF1A-TFEB-S141A/S210A-hygro, respectively), were custom-designed and packaged by VectorBuilder (Chicago, IL, USA). Detailed vector information can be accessed using the following IDs on vectorbuilder.com: VB241111-1068mpb (TFEB) and VB241111-1066uxv (TFEB^S141A/S210A^). Plasmid assembly was validated through the whole plasmid sequencing service provided by Eurofins Genomics US (Louisville, KY, USA).

**Table 1.**
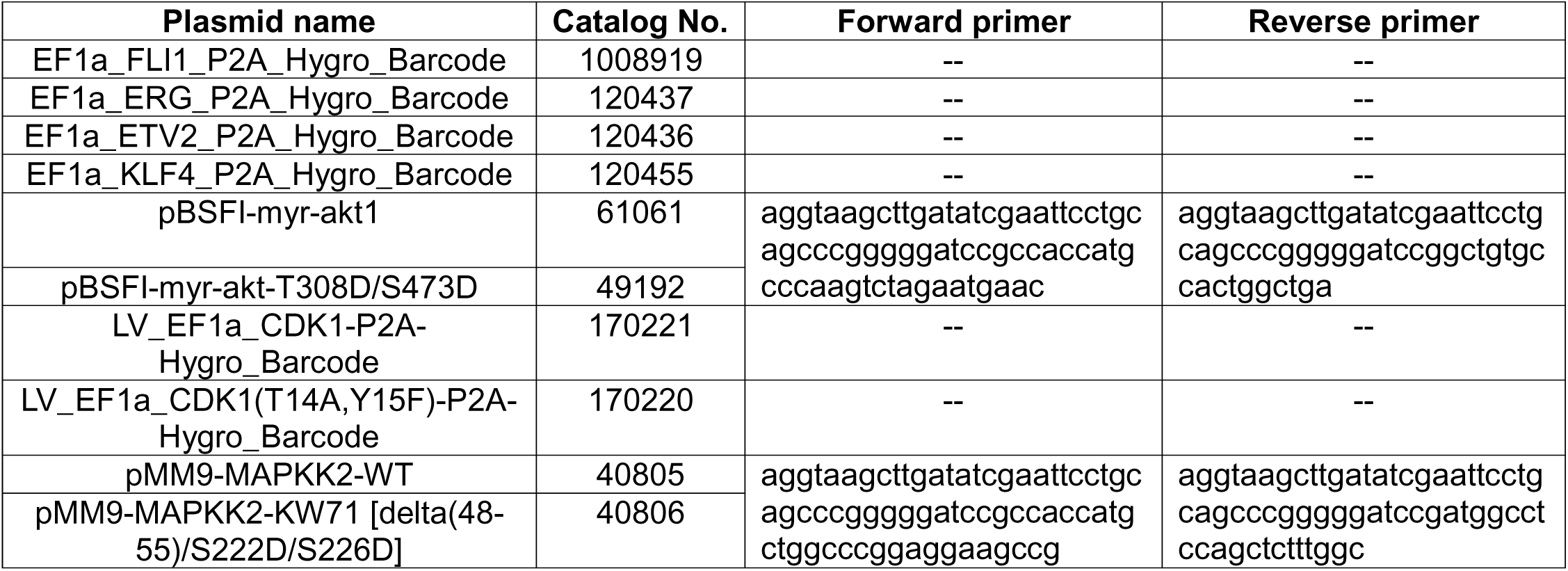
Plasmids and primers used for lentiviral particle assembly in this study.

### Lentiviral preparation

HEK293T cells (Cat. No. CRL-11268, ATCC, Manassas, VA, USA) were seeded at a density of 4 million cells per 10 cm dish and cultured overnight in DMEM (Cat. No. 11995065, Thermo Fisher Scientific, Waltham, MA, USA) supplemented with 10% FBS (Cat. No. CORMT35010CV, Corning, Corning, NY, USA). For transfection, a DNA mixture containing 5 μg of the gene of interest, 3 μg of PAX2, and 2 μg of MD2G was prepared in 600 μL of Opti-MEM (Cat. No. 31985079, Thermo Fisher Scientific, Waltham, MA, USA). Separately, 40 μL of Lipofectamine 2000 was mixed with 560 μL of Opti-MEM (Cat. No. 11668019, Thermo Fisher Scientific). After a 15-minute incubation, the two mixtures were combined and incubated for an additional 25 minutes. The final transfection mixture was then added dropwise to the HEK293T cells. At 48-and 60-hours post-transfection, the culture media containing lentiviral particles was collected and transferred to PET thin wall tubes (Cat. No. 75-000-471, Thermo Fisher Scientific). The lentivirus-containing media was centrifuged at 20,000 x g for 2 hours using a Sorvall WX80 Ultracentrifuge (Cat. No. 75000080, Thermo Fisher Scientific). After centrifugation, the supernatant was aspirated, and the viral pellet was resuspended in approximately 1 mL of DPBS (Cat. No. 14190144, Thermo Fisher Scientific), aliquoted into 20 μL portions, and stored at −80°C.

### Cell culture

MEFs were obtained from Lonza Bioscience (Cat. No. M-FB-481, Walkersville, MD, USA), and mouse aortic endothelial cells (MAOECs) were purchased from iXCells Biotechnologies (Cat. No. 10MU-002, San Diego, CA, USA). All cell lines were cultured in DMEM supplemented with 10% fetal bovine serum under standard conditions.

### Lentiviral infection

Fresh aliquots of lentivirus were thawed prior to each infection, and the viral concentration was assessed using a qPCR lentivirus titer kit (Cat. No. LV900, Applied Biological Materials, Richmond, BC, Canada). A concentration of 1,000 lentiviral genomes per cell was empirically determined to be non-toxic while effectively overexpressing the genes of interest; thus, this concentration was used for all infections. Cells were infected in DMEM + 10% FBS supplemented with 8 μg/mL polybrene (Cat. No. TR-1003-G, MilliporeSigma, Burlington, MA, USA). For experiments involving multiple viruses, each lentivirus was administered at 1,000 lentiviral genomes per cell. Twenty-four hours after infection, the culture medium was replaced, and cells were maintained in DMEM supplemented with 10% FBS. Control cells were infected with either an mCherry-expressing hygromycin vector or an empty pLKO lentiviral puromycin vector (Addgene plasmid #10879; https://www.addgene.org/10879/; RRID) at a multiplicity of 1,000 lentiviral genomes per cell. At 72 hours post-infection, selection was initiated by adding hygromycin (50 μg/mL; Cat. No. 10687010, Thermo Fisher Scientific, Waltham, MA, USA) or puromycin (1 μg/mL; Cat. No. ILTA1113803, Thermo Fisher Scientific, Waltham, MA, USA) to the culture medium to enrich for successfully transduced cells.

### Quantitative real-time PCR analysis

Cells were seeded at a density of 20,000 cells per well in 24-well plates (Cat. No. 12-556-006, Thermo Fisher Scientific). Each experimental condition, representing a specific viral infection, was performed in triplicate, with three wells allocated per condition. At 9 days post-infection, total RNA was extracted from the cells using Trizol reagent (Cat. No. ILT15596018, Thermo Fisher Scientific) following the manufacturer’s protocol. RNA concentrations were quantified using a NanoDrop One spectrophotometer (Cat. No. ND-ONE-W, Thermo Fisher Scientific).

To synthesize cDNA, 1 μg of RNA per 20 μL reaction was reverse transcribed using the iScript cDNA synthesis kit (Cat. No. 1708891, Bio-Rad Laboratories, Hercules, CA, USA). The cDNA was then diluted to a final RNA concentration of 5 ng/μL by adding 180 μL of nuclease-free water per 20 μL cDNA reaction. Quantitative RT-PCR was performed in triplicate using 20 μL reactions with SYBR Green Master Mix (Cat. No. ILT15596018, Thermo Fisher Scientific). The qPCR was run on a QuantStudio 5 Real-Time PCR System (Cat. No. A34322, Thermo Fisher Scientific) using the primers listed in **Table 2**. Cycle threshold (Ct) values were collected for analysis, and gene expression levels were normalized against the GAPDH housekeeping gene.

**Table 2.**
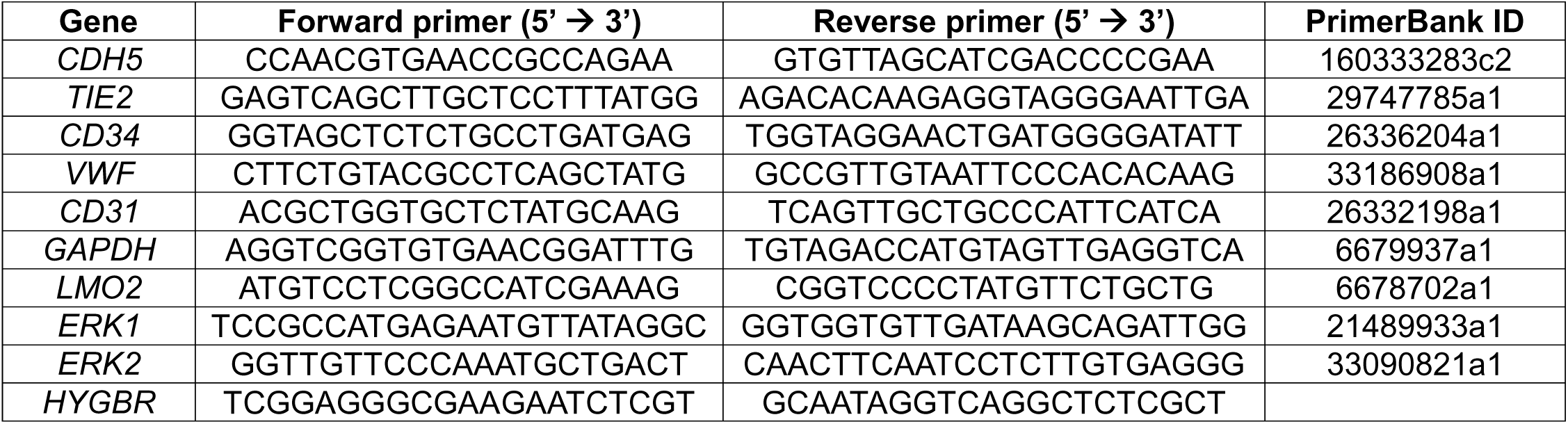
Primer sequences used for qRT-PCR analysis.

### Bulk RNA sequencing

Cells were seeded and infected under the same conditions described in the Quantitative real-time PCR analysis section. Total RNA was extracted using Trizol and stored in TE buffer (Cat. No. 12090015, Thermo Fisher Scientific). To enrich for mRNA, poly-A selection was performed prior to cDNA synthesis and library preparation using the xGen™ Broad-Range RNA Library Preparation Kit (Cat. No. 10009813, Integrated DNA Technologies, Coralville, IA, USA) and the NEB Poly(A) kit (Cat. No. E7490S, New England Biolabs) according to the manufacturers’ protocols. After poly-A enrichment, samples were pooled and subjected to a 151 bp paired-end sequencing run on a NovaSeq X Plus 10B Flow Cell (Illumina) per the manufacturer’s instructions. De-multiplexed Fastq files were generated using BCL Convert Conversion Software v4.0 (Illumina). Reads were trimmed with Cutadapt v2.33^19^ and quality was assessed using FastQC v0.11.8. Fastq Screen v0.13.0 was employed to detect contamination^20^, and reads were mapped to the reference genome GRCm38 (ENSEMBL), using STAR v2.7.8a.^21^ Gene-level counts were assigned using RSEM v1.3.3^22^, following ENCODE RNA-seq standards. Multiqc v1.7 was used to compile quality control metrics from these tools into a comprehensive report.^23^ Library prep and next-generation sequencing was carried out in the Advanced Genomics Core at the University of Michigan.

### Differential gene expression analysis

Differential gene expression analysis and heat map generation were performed using Python with the pyDESEq2 package.^24^ Genes with a log2 fold change greater than 2 and an adjusted p-value below 0.01 were considered differentially expressed. Adjusted p-values were calculated using the Benjamini & Hochberg (BH) method via the DeseqStats function in Python. Gene ontology (GO) enrichment analysis was conducted by selecting the top 250 most enriched genes, with GO terms determined using the Gene Ontology Resource (2024-09-08 release).^25–27^ Gene Set Enrichment Analysis (GSEA) was performed using the GSEApy package.^28^

### Lysosome staining

Lysosomes were labeled using a commercial lysosome-specific BioTracker dye (Cat. No. SCT138, Millipore Sigma). The dye was reconstituted in 100 μL of deionized water and applied to cells at a 1:500 dilution in culture medium. Fluorescence imaging was performed using a 620/60 nm filter cube on a Keyence BZ-X800 microscope (Cat. No. 26-1871514-64, Keyence, Osaka, Japan).

### Lysosome fluorescence measurement

Cells were seeded overnight at a density of 2,500 cells per well in clear glass-bottom, black-walled 384-well plates (Cat. No. 781097, Greiner Bio-One, Kremsmünster, Austria) using a Multidrop Combi dispenser (Cat. No. 5840300, Thermo Fisher Scientific, Waltham, MA, USA) with 20 μL of phenol red–free DMEM supplemented with 2% FBS (Cat. No. 31053028, Thermo Fisher Scientific). The following day, cells were either transduced with lentivirus using a multichannel pipette or treated with pharmacological compounds dispensed via an Echo 650 acoustic dispenser (Cat. No. 001-16079, Beckman Coulter, Brea, CA, USA). After compound and/or virus addition, an additional 20 μL of medium containing the lysosome-specific BioTracker dye (1:500 final dilution) was gently added using the Combi dispenser set to a medium-speed dispense mode to avoid cell disturbance. Lysosomal fluorescence intensity was measured using the PHERAstar FS microplate reader (Cat. No. KBS-0021-001, BMG LABTECH, Ortenberg, Germany) at 24 hours post–drug treatment or 48 hours post–lentiviral transduction.

### Mouse hindlimb ischemia

All animal procedures were approved by the University of Michigan Institutional Animal Care and Use Committee (IACUC). Hindlimb ischemia surgeries were performed on male C57BL/6J mice (Cat. No. 000664, The Jackson Laboratory, Farmington, CT, USA) between 6 and 10 weeks of age.^29,30^ Prior to surgery, the hindlimbs were shaved, and unilateral femoral artery ligation was performed on the left leg. Two sutures were placed approximately 3 mm apart to ligate the femoral artery distal to the origin of both the profunda femoris (deep femoral artery) and the superficial epigastric artery.

For cell transplantation experiments, MEFs were seeded at a density of 500,000 cells per 6-cm dish and transduced with either CA-MEK2 and FLI1 or mCherry control vectors for 9 days. On day 9, cells from each dish were harvested, suspended in 100 μL of sterile PBS, and injected intramuscularly into the ischemic hindlimb adjacent to the ligation site. Immediately after cell injection, the ligated 3-mm section of the femoral artery was surgically transected to induce complete ischemia. The surgical site was then sutured, and mice were allowed to recover. Analgesia was provided by subcutaneous injection of carprofen (0.5 mg/kg; Cat. No. 33975-100MG-R, MilliporeSigma) once daily for 72 hours postoperatively.

Laser Doppler perfusion imaging was performed using the PeriScan PIM 3 system (Cat. No. 44-00113-03, Perimed AB, Stockholm, Sweden) immediately after surgery (day 0) and again on day 7. Blood flow in the ischemic limb was normalized to the contralateral, uninjured limb using Perimed PIM 3 analysis software. During imaging sessions, body temperature was maintained at 37°C. Functional recovery of the hindlimb was also assessed based on clinical scoring of tissue damage.^31^

### In vitro angiogenesis

MEFs and MAOEC were seeded at a density of 500,000 cells per 6-cm dish and infected with 1,000 lentiviral genomes per cell to generate CA-MEK2 cells. Control MEFs were plated at the same density and infected with an equivalent titer of pLKO lentivirus. Cells were maintained for nine days post-infection before passaging and seeding onto Geltrex (Cat. No. A1413202, Thermo Fisher Scientific). On day 8 of transduction, pharmacologic treatments were initiated with 1 nM BafA1 (Cat. No. 11038, Cayman Chemical, Ann Arbor, MI, USA) or 1 nM CCA (Cat. No. 11050, Cayman Chemical). At the same time, Geltrex was transferred from −80 °C to 4 °C to thaw overnight.

On day 8, a 70 μL layer of Geltrex was dispensed into each well of a black-walled, clear glass-bottom 96-well plate (Cat. No. 655077, Greiner Bio-One). Pipette tips were pre-chilled on ice for 1 hour, and the plate was kept on ice during coating to prevent premature gelation. After dispensing, wells were inspected to ensure complete coverage and absence of bubbles, and the plate was then incubated at 37 °C for 1 hour to allow Geltrex polymerization. Cells were subsequently passaged and seeded at 20,000 cells per well and cultured overnight to allow angiogenic network formation. Cells treated with BafA1 or CCA were maintained in 1 nM of the respective compound during seeding in either DMEM + 10% FBS or endothelial growth media (EGM-2; Cat. No. MD-0010-500mL, iXCells Biotechnology, San Diego, CA, USA). Tube formation was assessed by live imaging the following day.

### Cell viability assay

Cells were seeded at a density of 10,000 cells per well in white-walled, white-bottom 96-well plates (Cat. No. 6055680, Revvity, Waltham, MA, USA). After overnight attachment, cells were treated with pharmacological compounds for 48 hours. Cell viability was then assessed using the CellTiter-Glo Luminescent Cell Viability Assay (Cat. No. G7572, Promega, Madison, WI, USA), following the manufacturer’s instructions. Luminescence was measured using a GloMax Discover Microplate Reader (Cat. No. GM3000, Promega).

### Small molecules

KO-947 (Cat. No. HY-112181), MK-8353 (Cat. No. HY-111407), EN6 (Cat. No. 1808714-73-9), and Torin-1 (Cat. No. HY-13003) were obtained from MedChemExpress (Monmouth, NJ, USA). BafA1 and CCA (Cat. No. 11038) or 1 nM CCA (Cat. No. 11050) were obtained from Cayman Chemical. All compounds were dissolved in DMSO to prepare 50 mM stock solutions.

### Immunofluorescence

Cells were seeded at a density of 10,000 cells per well in black-walled, clear glass-bottom 96-well plates (Cat. No. 655077, Greiner Bio-One) and transduced with lentivirus. On day 3 post-transduction, hygromycin (50 μg/mL) and pharmacological compounds were added to the culture medium. On day 5, cells were fixed in 3.6% paraformaldehyde (Cat. No. J19943.K2, Thermo Fisher Scientific) containing 0.1% Triton X-100 (Cat. No. HFH10, Thermo Fisher Scientific) for 15 minutes at room temperature, followed by three 5-minute washes with PBS.

Cells were then blocked in 4% heat-inactivated horse serum (Cat. No. 26050088, Thermo Fisher Scientific) prepared in PBS for 1 hour. After blocking, cells were incubated overnight at 4°C with a 1:100 dilution of anti-TFEB antibody (Cat. No. PA5-96632, Thermo Fisher Scientific) or phospho-p44/42 ERK1/2 (Cat. No. 9101S, Cell Signaling Technology, Danvers, MA, USA) in blocking solution. The following day, cells were washed three times with PBS (5 minutes each) and incubated for 2 hours at room temperature with Alexa Fluor 594 secondary antibody (1:200; Cat. No. A-21209, Thermo Fisher Scientific) and DAPI (0.1 μg/mL; Cat. No. D9542-1MG, Millipore Sigma, Burlington, MA, USA). After staining, cells were washed three additional times with DPBS and imaged directly in DPBS using a Keyence BZ-X800 fluorescence microscope. All washes and incubations were performed in 200 μL total volume per well.

### TFEB nuclear accumulation calculation

TFEB nuclear localization was quantified using the ROI Manager in FIJI (ImageJ) software.^32^ For each condition, five representative fluorescence images containing at least five cells each were analyzed. Regions of interest (ROIs) corresponding to individual nuclei were defined based on DAPI staining, and cytoplasmic ROIs were defined by the remaining cell area. The integrated fluorescence intensity of TFEB within nuclear ROIs (overlapping with DAPI) and the total cellular TFEB fluorescence were measured. The nuclear TFEB fraction was calculated as the integrated density of TFEB signal within DAPI-positive regions divided by the total TFEB integrated density for each cell. Mean nuclear TFEB ratios were then averaged across all cells and images per condition and statistical significance was determined by two-way ANOVA.

## Results

### Over-expression of constitutively active MEK2 (CA-MEK2) converts MEFs into the EC lineage

Post-translational modifications (PTMs) are central regulators of cell fate decisions, enabling rapid and reversible control of protein function, localization, and stability. Among these, phosphorylation is a dominant signaling modality that integrates extracellular cues with dynamic transcriptional programs during lineage transitions. To investigate how phosphorylation-driven signaling shapes endothelial reprogramming, we focused on three kinases—AKT1, CDK1, and MEK2—based on their well-established roles in phosphorylating key substrates that coordinate PTM networks governing cell identity and transcriptional plasticity.^33–36^ To systematically assess their reprogramming potential, we constructed six lentiviral vectors encoding either wild-type (WT) or constitutively active (CA) forms of AKT1, CDK1, and MEK2. Each construct was cloned into an identical lentiviral backbone containing a P2A-hygromycin resistance cassette to allow for consistent selection and expression across conditions. MEFs were transduced with these constructs and cultured for nine days in DMEM (Figure 1A). Successful transduction was confirmed by hygromycin resistance, and RT-qPCR analysis of hygromycin expression confirmed comparable transduction efficiency among all constructs (Figure S1A). To assess endothelial induction, we measured expression of canonical endothelial markers *(CDH5, CD31, CD34, VWF)* and the endothelial transcription factor *LMO2*. Among the tested kinases, CA-MEK2 markedly upregulated all five markers compared to mCherry controls (Figure 1B), whereas WT or CA forms of AKT1 and CDK1 showed minimal effects. These results suggested that sustained MEK2 activation may promote fibroblast reprogramming toward an endothelial-like phenotype.

**Figure 1:**
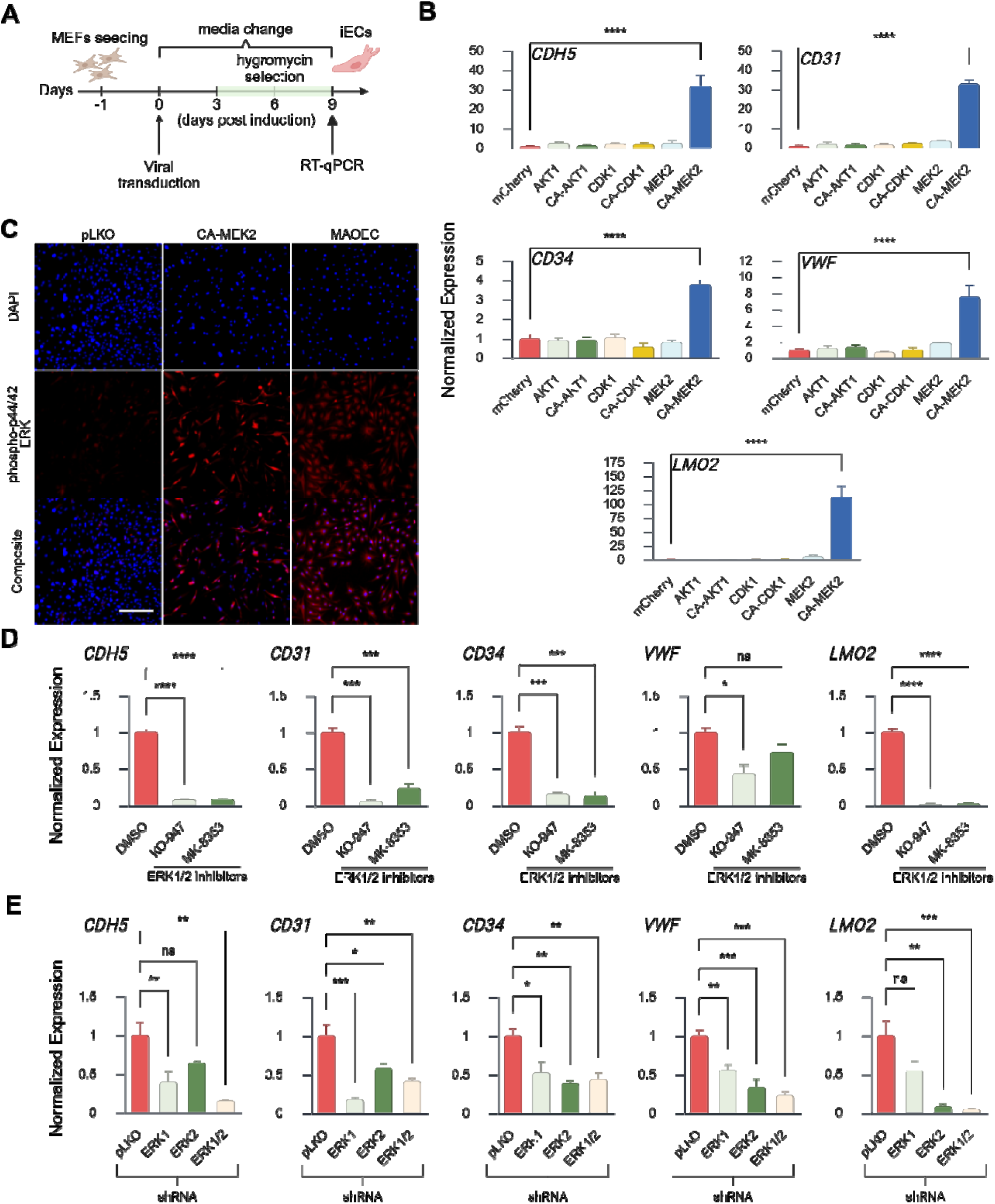
Over-expression of CA-MEK2 converts MEFs into the EC lineage. (A) Schematic diagram of the kinase screening strategy. Wildtype and constitutively active (CA) individual genes of kinases were transduced into MEFs. Medium was changed every 2 days with selection starting on day 3 post-transduction. Reprogramming efficiency was evaluated 9 days after transduction by qPCR. (B) Quantitative RT-PCR analysis showing relative expression of endothelial lineage genes (*CDH5, CD31, CD34, VWF,* and *LMO2*) in MEFs transduced with wild-type or CA kinases AKT1, CDK1, and MEK2. \*\*\*\**p* < 0.0001 vs. mCherry. N=3. (C) Immunofluorescence of phospho-p44/42 ERK1/2, downstream targets of CA-MEK2, expression (red) with DAPI nuclear counter stain (blue). Scale bar, 100 μm. (D) Quantitative RT-PCR analysis showing relative expression of endothelial lineage genes (*CDH5*, *CD31*, *CD34, VWF,* and *LMO2*) in CA-MEK2 transduced MEFs following treatment with ERK1/2 inhibitors KO-947 (1 μM) or MK-8353 (1 μM). ∗*p* < 0.05, \*\**p* < 0.01, \*\*\**p* < 0.001, and \*\*\*\**p* < 0.0001 vs. CA-MEK2 treatment with DMSO. N = 3. (E) Quantitative RT-PCR analysis showing relative expression of endothelial lineage genes (*CDH5*, *CD31*, *CD34, VWF,* and *LMO2*) in CA-MEK2 transduced MEFs in response to ERK1/2 shRNA knockdown. ∗*p* < 0.05, \*\**p* < 0.01, \*\*\**p* < 0.001, and \*\*\*\**p* < 0.0001 vs. CA-MEK2 treatment with empty pLKO shRNA. N = 3. Data represent three independent experiments and are presented as mean ± SEM. Each independent experiment consists of at least three technical replicates. Two-way ANOVA.

Given that MEK2 primarily phosphorylates and activates ERK1/2, we next evaluated ERK phosphorylation following CA-MEK2 transduction using immunofluorescence staining for phospho-p44/42 ERK1/2. As expected, CA-MEK2 expression led to robust ERK1/2 phosphorylation relative to control MEFs (Figure 1C). Interestingly, MAOECs also exhibited strong phospho-ERK1/2 signals, whereas control MEFs did not, indicating that sustained ERK1/2 activation may be an intrinsic feature of endothelial identity. To test whether ERK1/2 activation is required for CA-MEK2-mediated reprogramming, we inhibited ERK signaling using two pharmacologic ERK inhibitors (KO-947 and MK-8353) as well as shRNA-mediated knockdown of ERK1/2. Both approaches effectively suppressed ERK phosphorylation (Figure S1B, C). Inhibition or knockdown of ERK1/2 abrogated the CA-MEK2-induced upregulation of endothelial markers (Figure 1D, E), confirming that the reprogramming effect of CA-MEK2 depends on ERK1/2 signaling activity.

Because ERK1/2 activation appeared critical for endothelial identity, we next asked whether it could synergize with transcription factor–based reprogramming strategies. We generated three lentiviral constructs expressing the endothelial transcription factors ETV2, ERG, and FLI1, each in the same vector backbone as the kinase library. Although each factor was expressed at comparable levels in MEFs (Figure S2A) and induced limited upregulation of endothelial markers, including CDH5, CD31, CD34, VWF, and LMO2 (Figure S2B), these effects were markedly weaker than those driven by ERK1/2 activation alone (Figure S2C—E). When co-expressed with CA-MEK2, each transcription factor showed enhanced induction of endothelial markers (Figure S2C—E), suggesting some cooperative signaling between the ERK1/2 pathway and endothelial transcriptional networks. Conversely, pharmacologic inhibition of ERK1/2 with KO-947 or MK-8353 during transcription factor–driven reprogramming significantly reduced endothelial marker expression (Figure S2F—H). Collectively, these findings indicate that MEK2-ERK1/2 activation is a much more dominate driving force than examined endothelial lineage transcription factors for fibroblast-to-endothelial reprogramming, underscoring ERK1/2 as a key regulator of endothelial cell identity and reprogramming fidelity.

To validate the angiogenic properties of the iECs generated through our reprogramming strategy, we employed a well-established mouse hindlimb ischemia model. Based on the initial quantitative RT-PCR in vitro data demonstrating the highest induction of endothelial marker gene expression, we selected the combination CA-MEK2 and FLI1 for further testing. Hindlimb ischemia was induced in male C57BL/6J mice by ligating and transecting the femoral artery in the left hindlimb, distal to the origin of the profunda femoris and superficial epigastric arteries (Figure 2A). Immediately following induction of ischemia, the reprogrammed MEFs (CA-MEK2 + FLI1) were injected into the muscle tissue adjacent to the site of ligation. Laser Doppler imaging revealed that mice treated with CA-MEK2 + FLI1 iECs exhibited significantly enhanced perfusion of the ischemic hindlimb by day 7 post-surgery compared to mCherry-transduced control mice (Figure 2B, C). These improvements were not only quantitative but also evident upon gross visual inspection. Mice in the CA-MEK2 + FLI1 group showed markedly reduced signs of necrosis and tissue loss in the foot and digits, as confirmed by clinical grading of limb damage (Figure 2D, E). These results suggest that fibroblasts reprogrammed into iECs can contribute to vascular regeneration and functional recovery in ischemic tissue.

**Figure 2:**
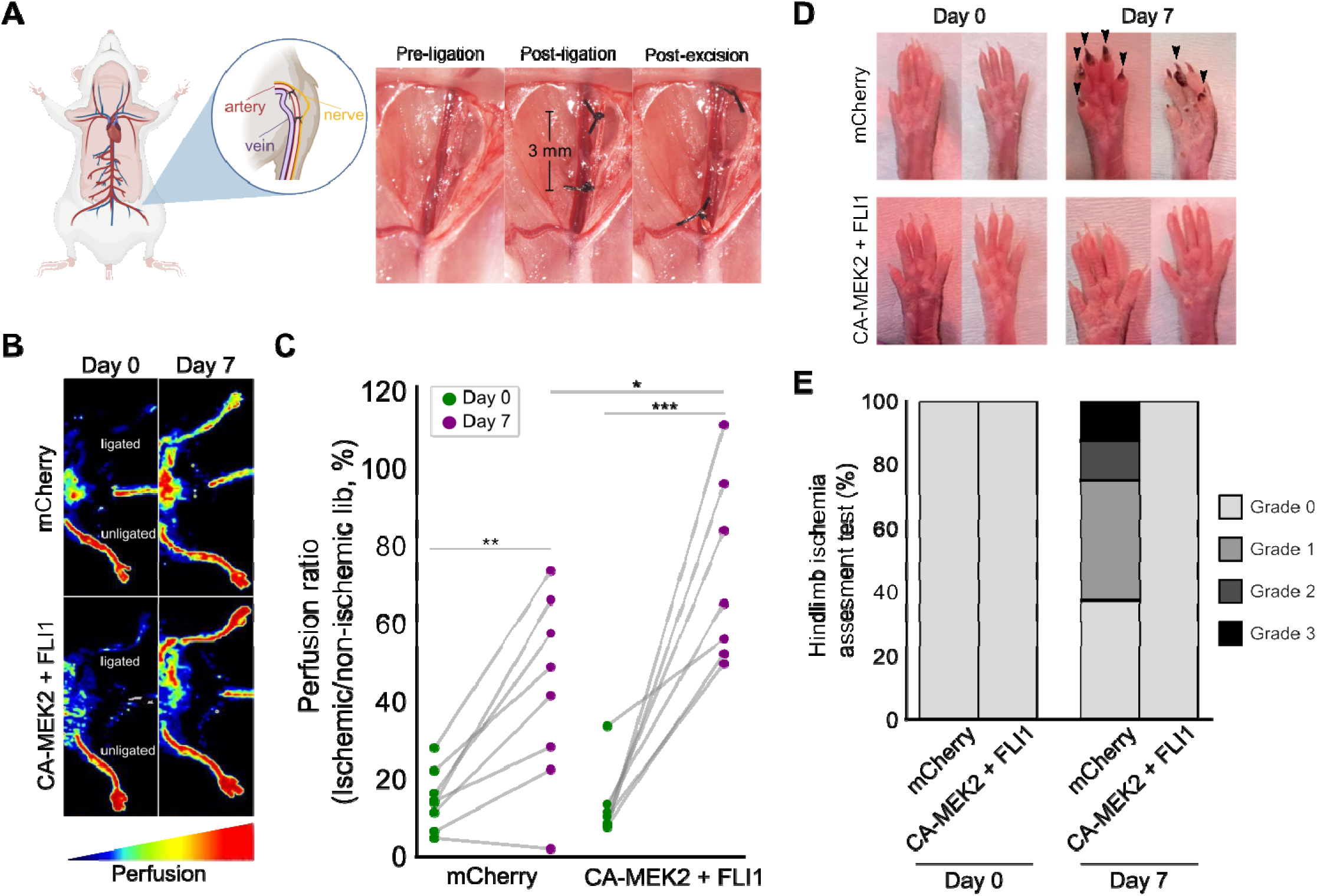
Induced endothelial cells promote functional recovery in ischemic hindlimb model. (A) Schematic representation of the femoral artery ligation procedure in the mouse hindlimb ischemia model. (B) Representative laser Doppler perfusion images (PeriScan PIM3 system) of ischemic limbs treated with mCherry control or CA-MEK2 + FLI1–transduced MEFs. (C) Quantification of perfusion ratio (ischemic limb/non-ischemic limb) at day 0 and day 7 post-surgery. (D) Representative photographs of distal hindlimb tissue showing gross morphology of ischemic limbs from each treatment group. (E) Clinical grading of ischemic limb damage based on a standardized scoring system as described by Wang et al., 2019.

### CA-MEK2 drives physical remodeling of the ECM, V-ATPase up-regulation, and lysosome acidification to facilitate endothelial reprogramming

To elucidate the transcriptional programs underlying CA-MEK2–driven endothelial reprogramming, we performed bulk RNA-seq on mCherry-control MEFs, FLI1-transduced MEFs, CA-MEK2–transduced MEFs, CA-MEK2 + FLI1 co-transduced MEFs, and mouse aortic endothelial cells (MAOECs) as a positive control. Principal component analysis (PCA) revealed distinct clustering of mCherry- and FLI1-transduced MEFs, as well as MAOECs, whereas CA-MEK2–transduced MEFs clustered closely with CA-MEK2 + FLI1 co-transduced MEFs, indicating a high degree of transcriptional similarity between these populations (Figure 3A). Notably, CA-MEK2 expression accounted for the predominant shift in cell identity, whereas FLI1 alone induced only modest transcriptional changes and did not substantially reposition MEFs toward the endothelial state. Based on this close relationship, we focused subsequent differential expression analyses on comparisons between mCherry-control MEFs and CA-MEK2–transduced MEFs.

**Figure 3:**
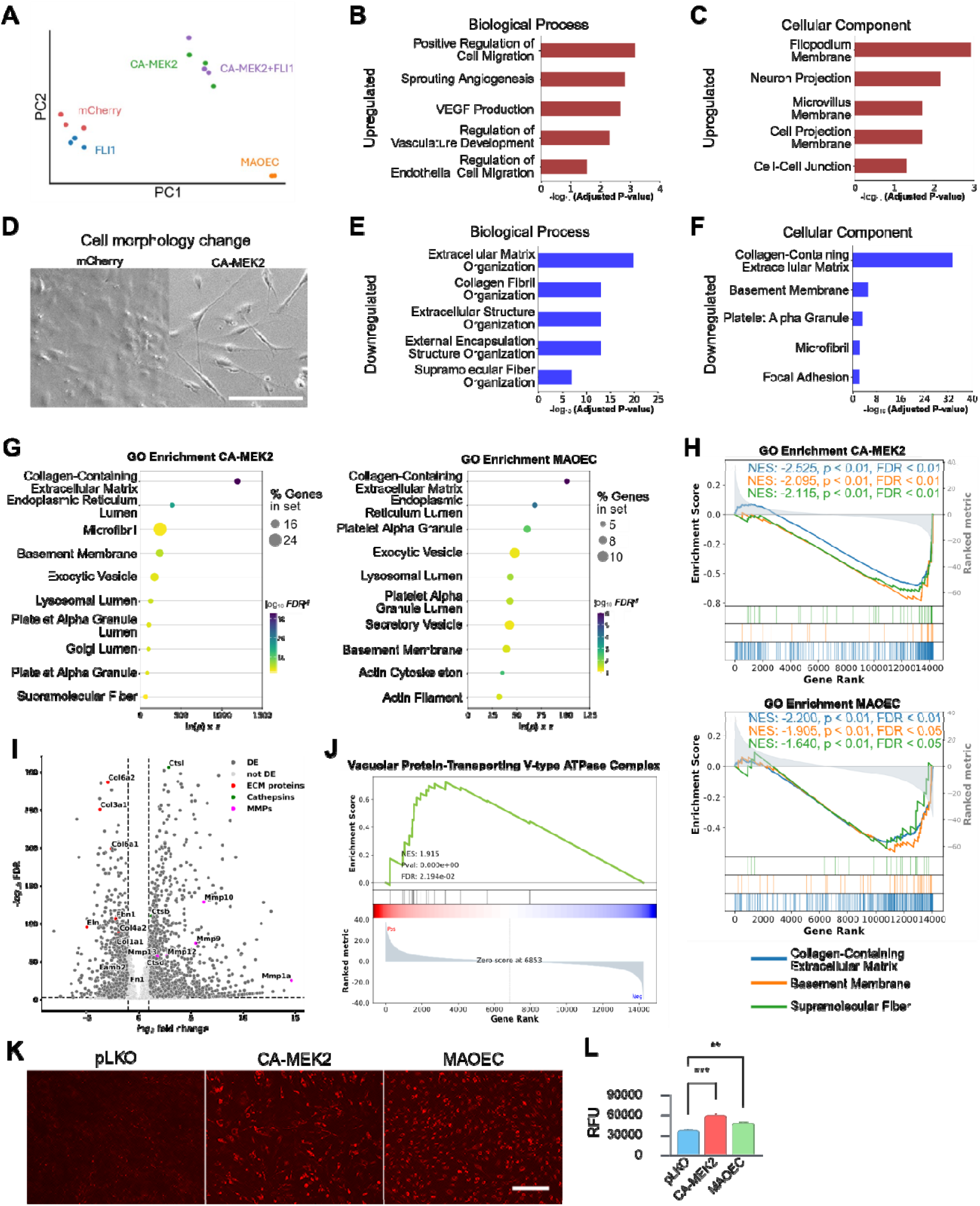
CA-MEK2 drives physical remodeling of the ECM, V-ATPase up-regulation, and lysosome acidification to facilitate endothelial reprogramming. (A) PCA analysis of RNA-seq groups. Gene expression was compared to mCherry-transduced controls. All data represent three biological replicates (N = 3). (B and C) Differential expression analysis highlighting the (A) biological process and (B) cellular component of the top 250 upregulated genes in CA-MEK2 vs. mCherry control. (D) Brightfield images representing morphological changes associated with mCherry control and CA-MEK2–transduced MEFs. Scale bar, 10Lμm. (E and F) Differential expression analysis highlighting the (D) biological process and (E) cellular component of the top 250 downregulated genes in CA-MEK2 vs. mCherry control. (G) GO analysis of downregulated differentially expressed genes in CA-MEK2–transduced MEFs (day 9) and MAOECs relative to mCherry controls. Dot size reflects the proportion of genes enriched within each GO cellular component category. (H) GSEA of extracellular matrix–related gene sets from the GO cellular component library in CA-MEK2–transduced MEFs and MAOECs versus mCherry controls. Highlighted gene sets include collagen-containing extracellular matrix, basement membrane, and supramolecular fiber. (I) Volcano plot of differentially expressed genes highlighting downregulation of ECM-related genes *(Col1a1, Col3a1, Col4a2, Col6a1, Col6a2, Eln, Fn1, Lamb2)* and upregulation of cathepsins *(Ctsb, Ctsl, Ctsd)* and metalloproteinases *(Mmp1a, Mmp9, Mmp10, Mmp12, Mmp13)* for CA-MEK2-transduced MEFs vs. mCherry controls. (J) GSEA of V-ATPase subunit genes from the Gene Ontology cellular component library comparing CA-MEK2–transduced MEFs to mCherry-transduced MEFs. (K) Representative fluorescence images of BioTracker NIR633 lysosome dye in pLKO-transduced, CA-MEK2–transduced MEFs, and MAOECs. Increased fluorescence intensity reflects enhanced lysosomal acidification. Scale bar, 100Lμm. (L) Quantification of lysosomal acidification using BioTracker NIR633 fluorescence in pLKO control, CA-MEK2–transduced MEFs, and MAOECs. Fluorescence intensity measured by PHERAstar plate reader (N = 16 wells per group). Data are presented as mean ± SEM; statistical significance assessed by two-way ANOVA.

Differential expression analysis between mCherry and CA-MEK2 conditions identified the top 250 up- and down-regulated genes. Gene ontology (GO) enrichment revealed that upregulated genes were strongly associated with endothelial biological processes, including angiogenesis, cell migration, and vasculature development (Figure 3B). Enrichment of cellular components related to the filopodium membrane (Figure 3C) was consistent with the observed morphological transition of CA-MEK2–transduced cells, which displayed elongated, branched morphologies resembling migratory endothelial cells (Figure 3D). To determine the lineage specificity of this conversion, we compared CA-MEK2 expression profiles with lineage-specific gene sets. Expression of 38 cardiomyocyte and skeletal muscle markers (Table S1) and 29 smooth muscle markers (Table S2) revealed that CA-MEK2–transduced MEFs clustered more closely with MAOECs than with mCherry controls (Figure S3A, B), confirming selective activation of endothelial, rather than general mesodermal, programs.

Analysis of downregulated genes revealed marked suppression of ECM–associated transcripts, particularly those encoding collagen-containing matrix components (Figure 3E, F). Because ECM remodeling is a defining feature of angiogenic endothelial cells, we performed gene set enrichment analysis (GSEA) across the full transcriptome, comparing CA-MEK2–expressing MEFs with MAOECs. The downregulated gene sets were strongly enriched for collagen fibril organization, basement membrane components, and supramolecular fibers (Figure 3G, H), indicating that CA-MEK2 activation promotes a shift toward an endothelial-like, matrix-remodeling phenotype.

To identify candidate effectors of this ECM remodeling, we examined the most significantly upregulated transcripts in CA-MEK2–transduced cells. Volcano plot analysis revealed increased expression of multiple cathepsins and matrix metalloproteinases (MMPs)—two proteolytic families responsible for ECM degradation (Figure 3I). Concurrently, several ECM structural genes appeared among the most strongly downregulated, consistent with enhanced ECM turnover.

Since cathepsins and MMPs require acidic conditions for proteolytic activation and ECM degradation^37–39^, a key function of CA-MEK2 could be to activate lysosome activity. Because lysosomal activation/acidification is carried out mainly by the V-ATPase proton pump, we next examined the expression of this pump. GSEA of the V-ATPase complex confirmed robust upregulation of multiple subunits of the V-ATPase in CA-MEK2–transduced cells relative to mCherry controls (Figure 3J). To test whether this transcriptional increase corresponded to functional acidification, we quantified lysosomal pH using a lysosome-specific pH-sensitive BioTracker dye. Both CA-MEK2–expressing MEFs and MAOECs displayed significantly greater lysosomal fluorescence intensity compared to control MEFs (Figure 3K, L), consistent with increased V-ATPase-dependent acidification.

Together, these findings identify sustained MEK2–ERK signaling as a potent and primary driver of fibroblast reprogramming toward an endothelial-like state, exerting effects that substantially exceed those achieved by endothelial transcription factors alone. Rather than acting as a secondary consequence, lysosomal activation—marked by enhanced V-ATPase–dependent acidification—emerges as a major downstream program engaged by MEK2–ERK signaling and closely associated with extracellular matrix remodeling. This coordinated increase in lysosomal function and ECM turnover likely underlies the acquisition of angiogenic behaviors, including migration and sprouting. Collectively, our transcriptomic and functional analyses position lysosomal acidification as a core downstream event in MEK2–ERK–driven endothelial reprogramming.

### V-ATPase functions as the major enzymatic driver of lysosome acidification and ECM remodeling promoting endothelial reprogramming

To test whether lysosomal acidification mediated by the V-ATPase complex plays a key role in endothelial reprogramming, we used pharmacological modulation of V-ATPase activity. We first applied the well-characterized V-ATPase inhibitors BafA1 and CCA to assess whether blocking lysosomal acidification would disrupt endothelial features. CA-MEK2–transduced MEFs and MAOECs were treated with 1 nM of either inhibitor for 24 h, followed by lysosomal pH imaging. As expected, both BafA1 and CCA markedly reduced fluorescence from the pH-sensitive lysosomal dye, confirming loss of organelle acidification in CA-MEK2–expressing and endothelial control cells (Figure 4A, B).

**Figure 4:**
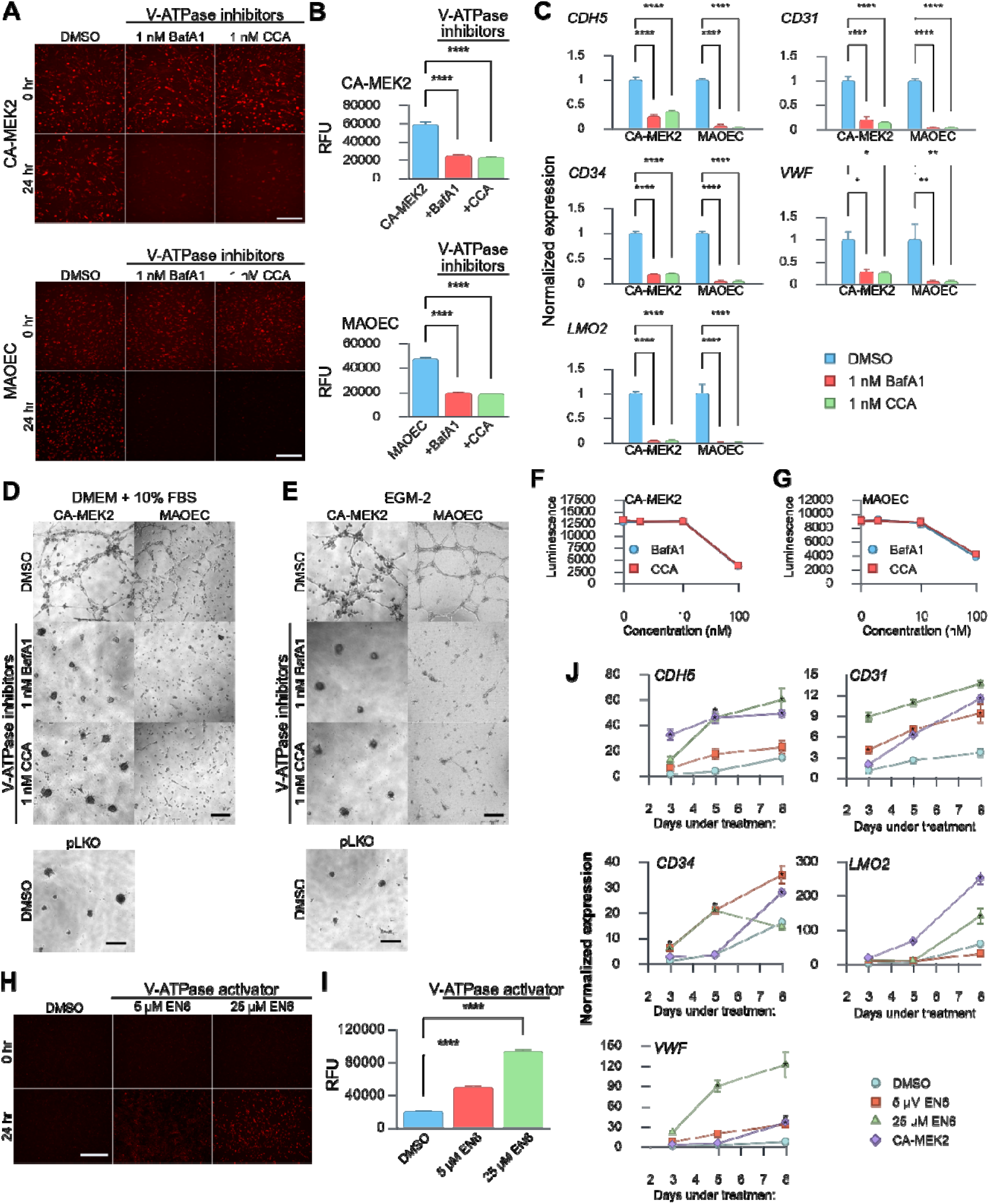
V-ATPase functions as the major enzymatic driver of lysosome acidification and ECM remodeling promoting endothelial reprogramming. (A) Representative fluorescence images of CA-MEK2–transduced MEFs and MAOECs stained with BioTracker NIR633 lysosome dye following treatment with DMSO, 1 nM bafilomycin A1 (BafA1), or 1 nM concanamycin A (CCA) for 0 or 24 hours. Scale bar, 100 μm. (B) Quantification of lysosomal acidification in CA-MEK2–transduced MEFs and MAOECs using BioTracker NIR633 fluorescence following 0 or 24 hours of incubation with DMSO, 1 nM BafA1, or 1 nM CCA. Fluorescence intensity was measured by PHERAstar plate reader (N = 16 wells per condition). (C) Quantitative RT-PCR analysis of endothelial marker genes *(CDH5, CD31, CD34, VWF,* and *LMO2)* in CA-MEK2–transduced MEFs and MAOECs after 24-hour treatment with DMSO, 1 nM BafA1, or 1 nM CCA. N = 3. *p < 0.05, **p < 0.01, ****p < 0.0001 versus DMSO control. (D and E) Representative brightfield images from in vitro tube formation assays using CA-MEK2–transduced MEFs and MAOECs following 24-hour treatment with DMSO, 1 nM BafA1, or 1 nM CCA. Cells were seeded in (D) DMEM + 10% FBS or (E) endothelial growth medium with added angiogenic factors. N = 3 per condition. Scale bar, 100 μm. (F and G) Cell viability measured by CellTiter-Glo luminescence in (F) CA-MEK2–transduced MEFs and (G) MAOECs after 48-hour exposure to DMSO, or 1 nM, 10 nM, or 100 nM BafA1 or CCA. (H) Representative fluorescence images of MEFs stained with BioTracker NIR633 lysosome dye after treatment with DMSO, 5 μM, or 25 μM EN6 for 0 or 24 hours. Scale bar, 100 μm. (I) Quantification of lysosomal acidification in MEFs after 24-hour treatment with DMSO, 5 μM, or 25 μM EN6. Fluorescence measured by PHERAstar plate reader (N = 16 wells per condition). (J) Quantitative RT-PCR analysis of endothelial marker genes *(CDH5, CD31, CD34, VWF,* and *LMO2)* in MEFs treated with DMSO, 5 μM, or 25 μM EN6 for 3, 5, and 8 days. N = 3. *p < 0.05 versus DMSO-treated controls. Data are presented as mean ± SEM; statistical significance determined by two-way ANOVA.

We next examined whether this loss of acidification altered the endothelial transcriptional program. CA-MEK2–transduced MEFs and MAOECs were incubated for 24 h with BafA1 or CCA, and expression of canonical endothelial markers *(CDH5, CD31, CD34, VWF, LMO2)* was quantified by RT-qPCR. Both inhibitors significantly decreased endothelial marker expression across cell types (Figure 4C), indicating that lysosomal acidification is essential for sustaining endothelial gene expression during fibroblast reprogramming and for maintaining endothelial identity in native ECs.

To evaluate the functional relevance of V-ATPase activity in angiogenic behavior, we performed in vitro tube-formation assays using Geltrex basement-membrane extract. In the absence of endothelial growth factors, CA-MEK2–transduced cells spontaneously organized into network-like tubular structures, whereas MAOECs, as expected, did not form tubes under these minimal conditions (Figure 4D). Addition of BafA1 or CCA completely abolished tube formation in CA-MEK2–expressing cells (Figure 4D), demonstrating that V-ATPase activity is necessary for this self-organized morphogenic behavior.

We next repeated the assay in endothelial growth medium supplemented with VEGFA and other pro-angiogenic factors. Under these enriched conditions, both CA-MEK2–transduced MEFs and MAOECs displayed robust tube formation (Figure 4E). However, inhibition of V-ATPase again prevented network assembly in both cell types, confirming that lysosomal acidification—and by extension, ECM remodeling—is indispensable for angiogenic morphogenesis. To ensure that the loss of tube formation was not due to cytotoxicity, we measured cellular ATP levels following 24-hour treatment with 1 nM BafA1 or CCA. Neither inhibitor significantly affected ATP production (Figure 4F, G), indicating that the observed inhibition of angiogenic behavior reflects a functional requirement for V-ATPase activity rather than reduced cell viability. Collectively, these findings identify V-ATPase as a critical effector of endothelial reprogramming and function.

To test whether V-ATPase activation alone could enhance endothelial reprogramming, we next employed EN6, a small-molecule V-ATPase agonist that covalently binds the ATP6V1A subunit to stimulate proton-pumping activity.^40^ Treatment of MEFs with EN6 (5 μM or 25 μM, 24 h) produced a strong increase in lysosomal acidification, as indicated by elevated BioTracker fluorescence (Figure 4H, I). We then assessed whether sustained lysosomal activation could induce endothelial gene expression. MEFs treated continuously with EN6 exhibited a progressive, time-dependent upregulation of *CDH5, CD31, CD34, VWF,* and *LMO2* over 3, 5, and 8 days (Figure 4J), indicating that V-ATPase activation alone can promote partial endothelial reprogramming.

Collectively, our findings establish a mechanistic link between sustained MEK2–ERK signaling and lysosomal activation, in which enhanced V-ATPase activity facilitates ECM degradation and supports angiogenic reprogramming. Moreover, direct pharmacologic activation of V-ATPase via EN6 reveals that stimulating lysosomal acidification alone can drive fibroblasts toward an endothelial-like transcriptional state, underscoring the lysosome’s role as a signaling hub in cell-fate reprogramming.

### V-ATPase activation drives TFEB nuclear translocation to potentially enhance endothelial reprogramming

Given that lysosomal acidification drives endothelial-like reprogramming of fibroblasts, we next sought to identify which components downstream of V-ATPase mediate this process. V-ATPase is a central hub for lysosomal signaling and interacts with multiple partners that influence cellular physiology, including mTORC1, which regulates growth by inhibiting V-ATPase activity; TFEB, which serves as a master transcriptional regulator of lysosomal biogenesis; and the Rag GTPases, which control V-ATPase complex assembly.^41–43^ EN6 activates V-ATPase by covalently binding the ATP6V1A subunit, leading to dissociation of mTORC1 from the lysosomal surface and allowing nuclear translocation of TFEB^40^. While these various partners could plausibly contribute to endothelial reprogramming, TFEB emerges as a particularly compelling candidate, as its transcriptional activity directly controls lysosomal gene networks and has been implicated in coordinating cellular remodeling.^44,45^ Further work has also validated that TFEB is a strong activator of angiogenesis, with endothelial-specific TFEB overexpression enhancing post-ischemic blood flow recovery, capillary density, vessel sprouting, and endothelial migration and tube formation, whereas TFEB deficiency impairs these responses in vitro and in vivo.^46^ We therefore examine if V-ATPase activation promotes endothelial-like reprogramming at least in part through TFEB-dependent transcriptional programs, positioning it as a logical focus for further mechanistic studies.

We examined TFEB localization under different conditions using immunofluorescence. In MEFs, endogenous TFEB was broadly distributed between the cytoplasm and nucleus (Figure 5A). Transduction with wild-type TFEB produced a similar pattern, confirming basal nucleocytoplasmic shuttling. However, expression of CA-MEK2 or a phospho-deficient TFEB mutant (TFEB^S141A/S210A^, which is resistant to mTORC1-mediated cytoplasmic retention) markedly increased TFEB nuclear accumulation (Figure 5A–B). Some residual cytoplasmic signal remained, suggesting partial but incomplete translocation under these conditions.

**Figure 5:**
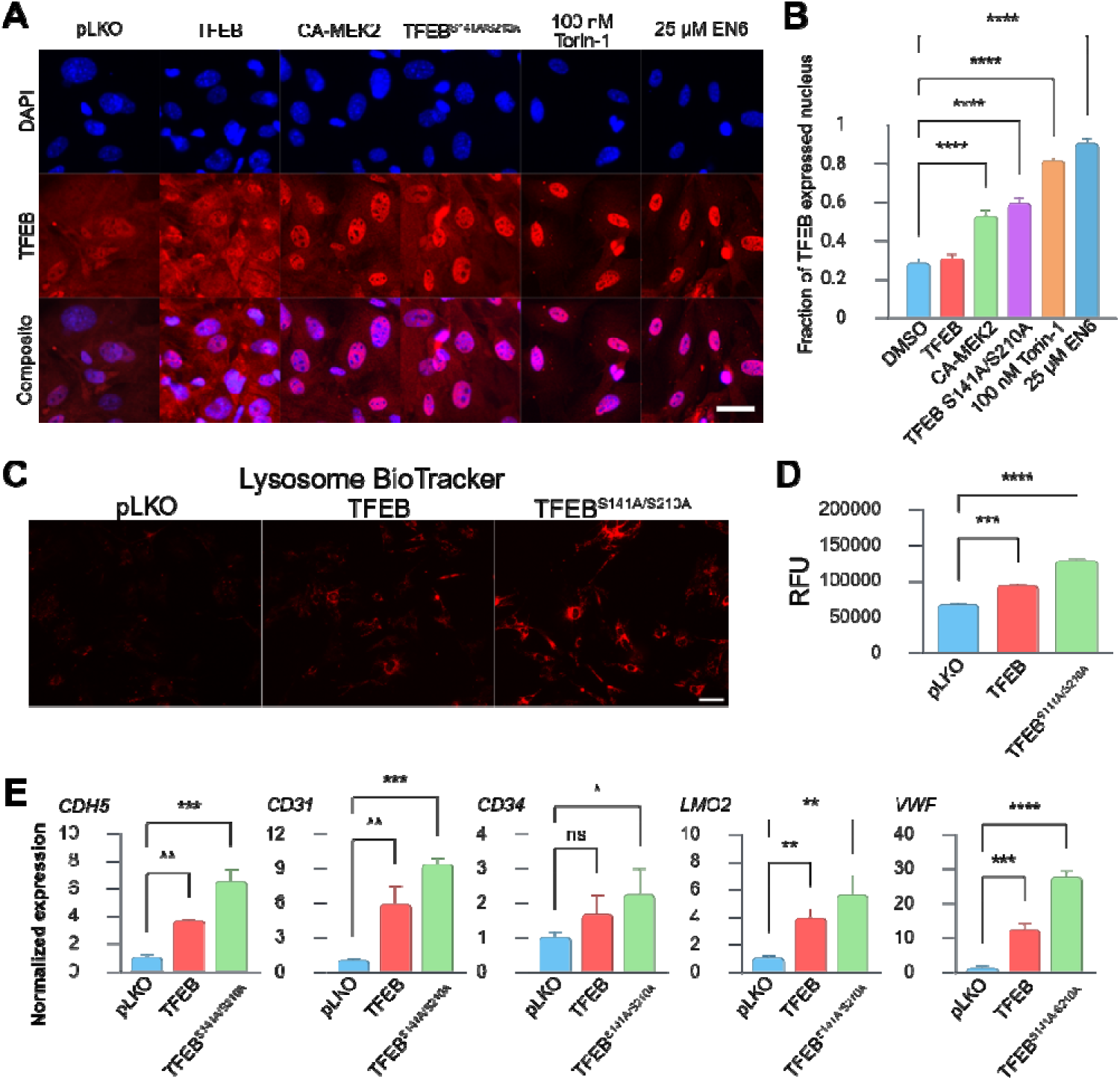
V-ATPase activation drives TFEB nuclear translocation to promote lysosome activity and endothelial gene expression. (A) Immunofluorescence images of MEFs transduced with genetic constructs (wildtype TFEB, CA-MEK2, or TFEB^S141A/S210A^), or treated with 100 nM Torin-1 or 25 μM EN6 for 2 days. TFEB staining is shown in red; nuclei are counterstained with DAPI (blue). Scale bar, 10 μm. (B) Quantification of TFEB nuclear localization based on integrated fluorescence intensity. Nuclear TFEB signal (within DAPI-positive regions) was divided by total cellular TFEB signal per cell. Nuclear ratios were averaged across ≥5 cells per image and across five independent experiments. Data are presented as mean ± SEM; statistical analysis by two-way ANOVA. (C) Representative fluorescence images of BioTracker NIR633 lysosome dye in MEFs transduced with pLKO control, wildtype TFEB, or TFEB^S141A/S210A^. Scale bar, 10 μm. (D) Quantification of lysosomal acidification using BioTracker NIR633 fluorescence in MEFs transduced with pLKO control, wild-type TFEB, or TFEB^S141A/S210A^. Fluorescence was measured using a PHERAstar plate reader (N = 16 wells per group). Data are shown as mean ± SEM; two-way ANOVA. (E) Quantitative RT-PCR analysis of endothelial marker genes *(CDH5, CD31, CD34, VWF,* and *LMO2)* in MEFs transduced with pLKO control, TFEB, or TFEB^S141A/S210A^. N=3. *p < 0.05, **p < 0.01, ***p < 0.001, ****p < 0.0001 vs. pLKO control; two-way ANOVA. Data are presented as mean ± SEM; statistical significance determined by two-way ANOVA.

Strikingly, stimulation with EN6 or Torin-1 (a potent mTOR inhibitor) led to near-complete TFEB nuclear translocation, consistent with full activation of the lysosome-to-nucleus signaling axis (Figure 5A, B). These findings indicate that V-ATPase activation—either through direct stimulation by EN6 or indirect mTOR inhibition—promotes TFEB nuclear localization.

We next evaluated whether TFEB activation enhances lysosomal function. MEFs transduced with wild-type TFEB or the S141A/S210A mutant exhibited significantly increased lysosomal fluorescence intensity compared with controls (Figure 5C, D), confirming that TFEB augments lysosomal activity. This supports the notion that V-ATPase activation triggers a positive feedback loop: lysosomal acidification promotes TFEB nuclear translocation, which in turn induces genes involved in lysosome biogenesis and function.

Finally, we asked whether TFEB activation alone is sufficient to induce an endothelial gene program. RT-qPCR analysis of TFEB and TFEB^S141A/S210A^ transduced MEFs revealed stepwise increases in the expression of endothelial markers *CDH5, CD31, CD34, VWF,* and *LMO2* (Figure 5E). While wild-type TFEB significantly upregulated these genes relative to controls, the nuclear-localized S141A/S210A mutant produced an even greater induction. These results indicate that the transcriptional activity of TFEB—particularly when localized to the nucleus—is capable of promoting endothelial gene expression in fibroblasts.

Collectively, these findings establish TFEB as a key downstream effector of V-ATPase-mediated reprogramming. Activation of V-ATPase through either sustained MEK2-ERK signaling or pharmacologic stimulation (EN6) enhances lysosomal acidification, releases mTORC1 inhibition, and drives TFEB nuclear translocation. Once activated, TFEB amplifies lysosomal and metabolic programs that facilitate ECM remodeling and angiogenic gene expression, reinforcing fibroblast conversion toward an endothelial-like phenotype.

## Discussion

In this study, we demonstrate that fibroblasts can be efficiently reprogrammed toward an endothelial-like state by harnessing lysosomal activity, rather than relying solely on transcription factor overexpression. Sustained MEK2–ERK signaling drives this conversion, engaging a MAPK–V-ATPase–TFEB axis that promotes lysosomal acidification, extracellular matrix remodeling, and angiogenic behavior. Pharmacologic or genetic modulation of lysosomal function further confirms its central role in shaping endothelial identity. Together, these findings establish lysosomes as an active regulator of cell fate and introduce an organelle-centric framework for reprogramming strategies that complements and, in some contexts, surpasses traditional transcription factor–based approaches.

These findings reframe ERK1/2 signaling in cell fate regulation by highlighting the importance of signaling duration. While ERK activity is classically associated with proliferation and reinforcement of mesenchymal identity ^47–49^, sustained ERK activation in our system produced a distinct outcome, transcriptionally upregulating multiple V-ATPase subunits and enhancing lysosomal acidification and proteolytic capacity. This lysosomal remodeling destabilized mesenchymal identity and redirected cells toward an endothelial-like state, indicating that ERK signaling is not required for mesenchymal fate maintenance. ERK-dependent activation of lysosomal function was accompanied by robust nuclear translocation of the lysosomal transcription factor TFEB, despite canonical models predicting enhanced cytosolic retention under sustained ERK signaling.^50–52^ We propose that strong ERK-driven V-ATPase upregulation induces a lysosomal stress–like state that impairs mTORC1 activity or its association with the lysosomal surface, thereby overriding ERK- and mTORC1-dependent sequestration of TFEB.^53^ Nuclear TFEB is then positioned to reinforce endothelial reprogramming through activation of gene programs controlling lysosomal biogenesis, autophagy, angiogenesis, cell migration, and extracellular matrix remodeling.^46,54–56^

Significantly, in contrast to transcription factor–centric reprogramming strategies, our approach targets the lysosome, a central regulator of cellular homeostasis, to induce broad phenotypic changes associated with endothelial identity. Traditional direct reprogramming relies on forced expression of lineage-defining transcription factors, such as ETV2, which can bias fibroblasts toward an endothelial fate but often with low efficiency and incomplete maturation.^8,9,57^ Although combinations of transcription factors modestly improve these outcomes, such approaches remain gene-centric and frequently require additional selection or culture manipulations to reinforce lineage commitment.^13^ Rather than imposing cell fate from forced expression of lineage specific transcription factors, our findings establish an organelle-targeted reprogramming strategy in which enhancing lysosomal function is sufficient to promote fibroblast conversion toward endothelial-like cells, accompanied by robust matrix remodeling and activation of endothelial gene programs (Figure 6). This framework represents a conceptual shift in reprogramming strategy, positioning organelle function as a central and instructive lever for cell fate control. By extension, we propose that once a cell acquires the defining structural and functional properties of a target lineage, here driven by lysosomal activation, it will behave as that lineage and progressively stabilize its identity over time.

**Figure 6:**
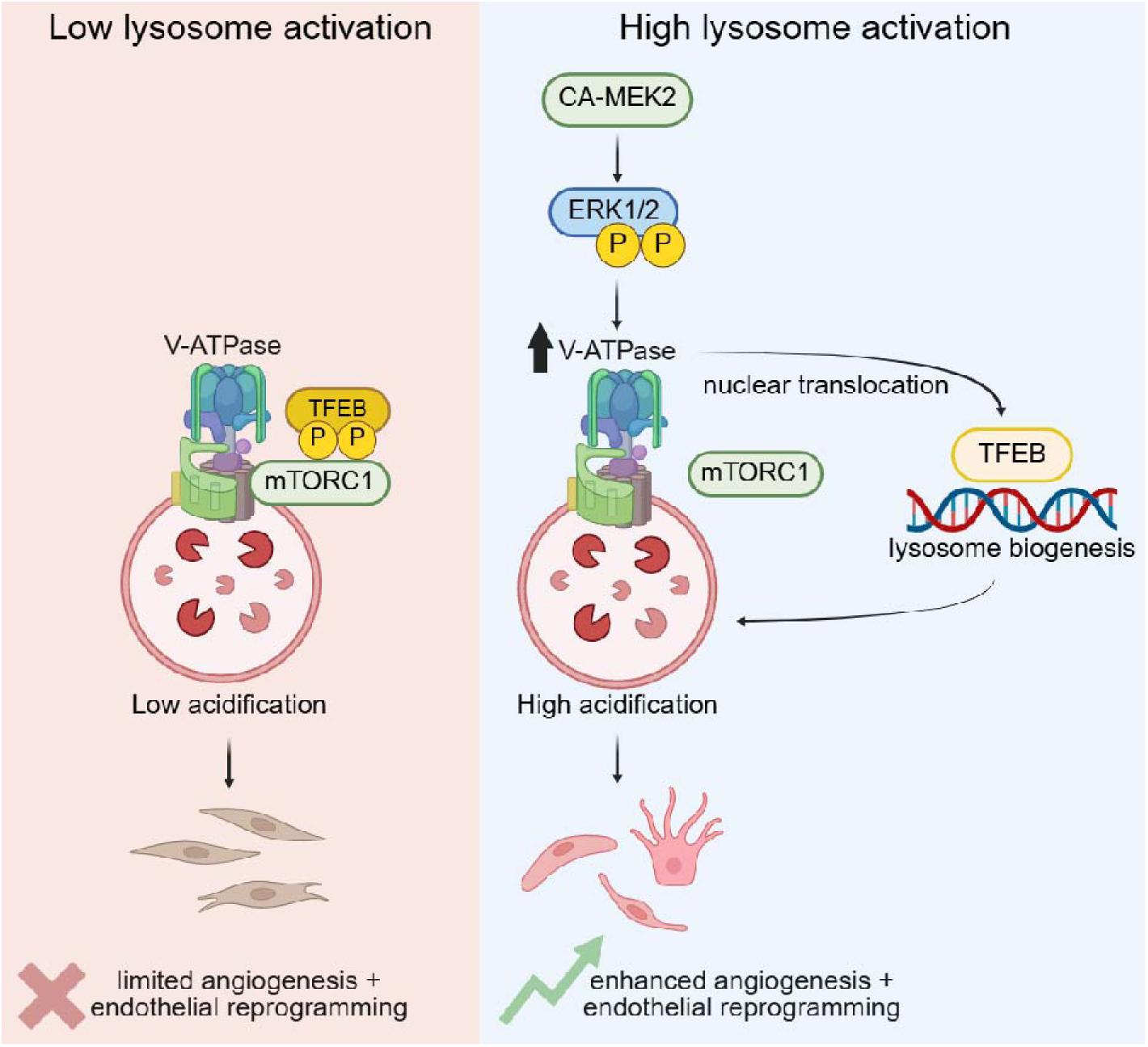
Central concept of lysosome mediated endothelial reprogramming. (Left) Reduced V-ATPase activity leads to impaired lysosomal acidification, constraining lysosome-dependent signaling and limiting angiogenesis and endothelial reprogramming. (Right) CA-MEK2 drives sustained ERK1/2 phosphorylation, which transcriptionally up-regulates V-ATPase subunits and enhances lysosomal acidification. Elevated lysosomal function promotes TFEB nuclear translocation, promoting lysosome biogenesis and contributing to angiogenesis and endothelial reprogramming.

This work is motivated by the principle that cell identity is ultimately defined by what a cell does, not merely by what it expresses. If a cell acquires the functional behaviors, structural organization, and environmental interactions of a target lineage, it should progressively adopt and stabilize the corresponding transcriptional state; thus, enforcing target cell properties provides a causal route to cell fate change. Our data support this framework by demonstrating that activation of lysosomal acidification induces hallmark endothelial behaviors, including ECM remodeling and angiogenic motility, and we therefore propose that driving these functional properties is sufficient to push fibroblasts across the endothelial identity threshold through an organelle-centric mechanism of transdifferentiation. In particular, our endothelial reprogramming approach uniquely leverages microenvironment remodeling: in fibrotic tissue, activated fibroblasts deposit abundant, stiff ECM that reinforces their identity^4–6^, yet by enhancing lysosomal acidification and protease activity, we enable these cells to degrade their surrounding matrix—mimicking the invasive behavior required for endothelial sprouting and migration.^58,59^ This functional remodeling of the microenvironment goes beyond simple activation of endothelial genes and anchors the transition toward a stabilized endothelial-like state.

In addition, these results point to a generalizable strategy in which identity erasure and target function promotion operate in parallel and potentially synergistic pathways. We propose that elevated lysosomal V-ATPase activity may broadly enhance reprogramming across lineages by accelerating the removal of residual features of the starting cell type. When coupled with interventions that promote hallmark functional activities of the target lineage, reprogramming efficiency may be further amplified. For example, in cardiomyocyte reprogramming, activation of lysosomal V-ATPase to erase the starting identity, together with strategies that promote contractile or electrophysiological activity, may synergistically reinforce acquisition of the cardiomyocyte state. Together, this framework emphasizes functional and organelle-level remodeling as an integral and actionable axis for improving cell fate reprogramming.

While our findings establish lysosomal activation as a key driver of endothelial reprogramming, several limitations warrant consideration. First, our work was performed entirely in vitro, leaving open the question of whether MEK2–V-ATPase–TFEB–driven reprogramming can be achieved efficiently and safely in vivo, where tissue context, immune interactions, and vascular architecture may influence outcomes. Second, V-ATPase has multifaceted roles in cellular physiology beyond lysosomal acidification, including regulation of pH homeostasis, vesicular trafficking, and metabolic signaling, and our study does not fully disentangle which aspects are most critical for endothelial identity. Finally, while we identified lysosomal activation as a permissive driver, the upstream cues and potential combinatorial factors that optimize reprogramming remain to be explored. Addressing these questions in future studies, through in vivo lineage tracing, conditional V-ATPase modulation, and combinatorial signaling interventions, will be essential for translating organelle-centric reprogramming strategies toward regenerative applications.

This study demonstrates that manipulating an organelle’s function can redirect cell fate, based on the principle that once a cell acquires the defining structural and functional properties of a target lineage, it will behave as that lineage and progressively stabilize its identity over time. From a therapeutic perspective, our fibroblast-to-endothelial conversion approach could be applied to promote cardiovascular repair after ischemic injury. The adult heart after myocardial infarction contains abundant activated fibroblasts and scar tissue but lacks sufficient vasculature. Converting some of those resident fibroblasts into endothelial cells in situ could help revascularize the injured region and mitigate fibrosis. Direct reprogramming approaches are particularly attractive here because they provide new cells for tissue repair while simultaneously reducing the fibrotic burden. Our organelle-based method could complement existing strategies (such as delivering pro-angiogenic factors or cells) by programmatically turning the scar tissue itself into blood vessels, leading to more integrated and long-lasting vessel formation within the infarcted myocardium.

## References

1. Cooley, B.C., Nevado, J., Mellad, J., Yang, D., St. Hilaire, C., Negro, A., Fang, F., Chen, G., San, H., Walts, A.D., et al. (2014). TGF-β Signaling Mediates Endothelial-to-Mesenchymal Transition (EndMT) During Vein Graft Remodeling. Sci. Transl. Med. 6. 10.1126/scitranslmed.3006927.

2. Qian, C., Dong, G., Yang, C., Zheng, W., Zhong, C., Shen, Q., Lu, Y., and Zhao, Y. (2025). Broadening horizons: molecular mechanisms and disease implications of endothelial-to-mesenchymal transition. Cell Communication and Signaling 23, 16. 10.1186/s12964-025-02028-y.

3. Alvandi, Z., and Bischoff, J. (2021). Endothelial-Mesenchymal Transition in Cardiovascular Disease. Arterioscler. Thromb. Vasc. Biol. 41, 2357–2369. 10.1161/ATVBAHA.121.313788.

4. Klingberg, F., Hinz, B., and White, E.S. (2013). The myofibroblast matrix: implications for tissue repair and fibrosis. J. Pathol. 229, 298–309. 10.1002/path.4104.

5. van den Borne, S.W.M., Diez, J., Blankesteijn, W.M., Verjans, J., Hofstra, L., and Narula, J. (2010). Myocardial remodeling after infarction: the role of myofibroblasts. Nat. Rev. Cardiol. 7, 30–37. 10.1038/nrcardio.2009.199.

6. Niro, F., Fernandes, S., Cassani, M., Apostolico, M., Oliver-De La Cruz, J., Pereira-Sousa, D., Pagliari, S., Vinarsky, V., Zdráhal, Z., Potesil, D., et al. (2024). Fibrotic extracellular matrix impacts cardiomyocyte phenotype and function in an iPSC-derived isogenic model of cardiac fibrosis. Translational Research 273, 58–77. 10.1016/j.trsl.2024.07.003.

7. Tani, H., Sadahiro, T., Yamada, Y., Isomi, M., Yamakawa, H., Fujita, R., Abe, Y., Akiyama, T., Nakano, K., Kuze, Y., et al. (2023). Direct Reprogramming Improves Cardiac Function and Reverses Fibrosis in Chronic Myocardial Infarction. Circulation 147, 223–238. 10.1161/CIRCULATIONAHA.121.058655.

8. Lee, S., Park, C., Han, J.W., Kim, J.Y., Cho, K., Kim, E.J., Kim, S., Lee, S.-J., Oh, S.Y., Tanaka, Y., et al. (2017). Direct Reprogramming of Human Dermal Fibroblasts Into Endothelial Cells Using ER71/ETV2. Circ. Res. 120, 848–861. 10.1161/CIRCRESAHA.116.309833.

9. Kim, J.-J., Kim, D.-H., Lee, J.Y., Lee, B.-C., Kang, I., Kook, M.G., Kong, D., Choi, S.W., Woo, H.-M., Kim, D.-I., et al. (2020). cAMP/EPAC Signaling Enables ETV2 to Induce Endothelial Cells with High Angiogenesis Potential. Molecular Therapy 28, 466–478. 10.1016/j.ymthe.2019.11.019.

10. Farber, G., Dong, Y., Wang, Q., Rathod, M., Wang, H., Dixit, M., Keepers, B., Xie, Y., Butz, K., Polacheck, W.J., et al. (2024). Direct conversion of cardiac fibroblasts into endothelial-like cells using Sox17 and Erg. Nat. Commun. 15, 4170. 10.1038/s41467-024-48354-6.

11. Ginsberg, M., James, D., Ding, B.-S., Nolan, D., Geng, F., Butler, J.M., Schachterle, W., Pulijaal, V.R., Mathew, S., Chasen, S.T., et al. (2012). Efficient Direct Reprogramming of Mature Amniotic Cells into Endothelial Cells by ETS Factors and TGFβ Suppression. Cell 151, 559–575. 10.1016/j.cell.2012.09.032.

12. Cho, S., Aakash, P., Lee, S., and Yoon, Y. (2023). Endothelial cell direct reprogramming: Past, present, and future. J. Mol. Cell. Cardiol. 180, 22–32. 10.1016/j.yjmcc.2023.04.006.

13. Grath, A., and Dai, G. (2024). SOX17/ETV2 improves the direct reprogramming of adult fibroblasts to endothelial cells. Cell Reports Methods 4, 100732. 10.1016/j.crmeth.2024.100732.

14. Prabakaran, S., Lippens, G., Steen, H., and Gunawardena, J. (2012). Post-translational modification: nature’s escape from genetic imprisonment and the basis for dynamic information encoding. Wiley Interdiscip. Rev. Syst. Biol. Med. 4, 565–583. 10.1002/wsbm.1185.

15. Powers, S.K., Holehouse, A.S., Korasick, D.A., Schreiber, K.H., Clark, N.M., Jing, H., Emenecker, R., Han, S., Tycksen, E., Hwang, I., et al. (2019). Nucleo-cytoplasmic Partitioning of ARF Proteins Controls Auxin Responses in Arabidopsis thaliana. Mol. Cell 76, 177–190.e5. 10.1016/j.molcel.2019.06.044.

16. Sabari, B.R., Dall’Agnese, A., Boija, A., Klein, I.A., Coffey, E.L., Shrinivas, K., Abraham, B.J., Hannett, N.M., Zamudio, A. V., Manteiga, J.C., et al. (2018). Coactivator condensation at super-enhancers links phase separation and gene control. Science (1979). 361. 10.1126/science.aar3958.

17. Liu, L., Lei, I., Tian, S., Gao, W., Guo, Y., Li, Z., Sabry, Z., Tang, P., Chen, Y.E., and Wang, Z. (2024). 14-3-3 binding motif phosphorylation disrupts Hdac4-organized condensates to stimulate cardiac reprogramming. Cell Rep. 43, 114054. 10.1016/j.celrep.2024.114054.

18. Parekh, U., Wu, Y., Zhao, D., Worlikar, A., Shah, N., Zhang, K., and Mali, P. (2018). Mapping Cellular Reprogramming via Pooled Overexpression Screens with Paired Fitness and Single-Cell RNA-Sequencing Readout. Cell Syst. 7, 548–555.e8. 10.1016/j.cels.2018.10.008.

19. Martin, M. (2011). Cutadapt removes adapter sequences from high-throughput sequencing reads. EMBnet. J. 17, 10. 10.14806/ej.17.1.200.

20. Wingett, S.W., and Andrews, S. (2018). FastQ Screen: A tool for multi-genome mapping and quality control. F1000Res. 7, 1338. 10.12688/f1000research.15931.2.

21. Dobin, A., Davis, C.A., Schlesinger, F., Drenkow, J., Zaleski, C., Jha, S., Batut, P., Chaisson, M., and Gingeras, T.R. (2013). STAR: ultrafast universal RNA-seq aligner. Bioinformatics 29, 15–21. 10.1093/bioinformatics/bts635.

22. Li, B., and Dewey, C.N. (2011). RSEM: accurate transcript quantification from RNA-Seq data with or without a reference genome. BMC Bioinformatics 12, 323. 10.1186/1471-2105-12-323.

23. Ewels, P., Magnusson, M., Lundin, S., and Käller, M. (2016). MultiQC: summarize analysis results for multiple tools and samples in a single report. Bioinformatics 32, 3047–3048. 10.1093/bioinformatics/btw354.

24. Muzellec, B., Teleńczuk, M., Cabeli, V., and Andreux, M. (2023). PyDESeq2: a python package for bulk RNA-seq differential expression analysis. Bioinformatics 39. 10.1093/bioinformatics/btad547.

25. Thomas, P.D., Ebert, D., Muruganujan, A., Mushayahama, T., Albou, L., and Mi, H. (2022). PANTHER: Making genome-scale phylogenetics accessible to all. Protein Science 31, 8–22. 10.1002/pro.4218.

26. Ashburner, M., Ball, C.A., Blake, J.A., Botstein, D., Butler, H., Cherry, J.M., Davis, A.P., Dolinski, K., Dwight, S.S., Eppig, J.T., et al. (2000). Gene Ontology: tool for the unification of biology. Nat. Genet. 25, 25–29. 10.1038/75556.

27. Aleksander, S.A., Balhoff, J., Carbon, S., Cherry, J.M., Drabkin, H.J., Ebert, D., Feuermann, M., Gaudet, P., Harris, N.L., Hill, D.P., et al. (2023). The Gene Ontology knowledgebase in 2023. Genetics 224. 10.1093/genetics/iyad031.

28. Fang, Z., Liu, X., and Peltz, G. (2023). GSEApy: a comprehensive package for performing gene set enrichment analysis in Python. Bioinformatics 39. 10.1093/bioinformatics/btac757.

29. Kim, H., Cho, H.-J., Kim, S.-W., Liu, B., Choi, Y.J., Lee, J., Sohn, Y.-D., Lee, M.-Y., Houge, M.A., and Yoon, Y. (2010). CD31+ Cells Represent Highly Angiogenic and Vasculogenic Cells in Bone Marrow. Circ. Res. 107, 602–614. 10.1161/CIRCRESAHA.110.218396.

30. Niiyama, H., Huang, N.F., Rollins, M.D., and Cooke, J.P. (2009). Murine Model of Hindlimb Ischemia. Journal of Visualized Experiments. 10.3791/1035-v.

31. Wang, L., Chen, Z., Li, Y., Yang, J., and Li, Y. (2019). Optical coherence tomography angiography for noninvasive evaluation of angiogenesis in a limb ischemia mouse model. Sci. Rep. 9, 5980. 10.1038/s41598-019-42520-3.

32. Schindelin, J., Arganda-Carreras, I., Frise, E., Kaynig, V., Longair, M., Pietzsch, T., Preibisch, S., Rueden, C., Saalfeld, S., Schmid, B., et al. (2012). Fiji: an open-source platform for biological-image analysis. Nat. Methods 9, 676–682. 10.1038/nmeth.2019.

33. Zhu, Z., Shukla, A., Ramezani-Rad, P., Apgar, J.R., and Rickert, R.C. (2019). The AKT isoforms 1 and 2 drive B cell fate decisions during the germinal center response. Life Sci. Alliance 2, e201900506. 10.26508/lsa.201900506.

34. Sakurikar, N., Eichhorn, J.M., and Chambers, T.C. (2012). Cyclin-dependent Kinase-1 (Cdk1)/Cyclin B1 Dictates Cell Fate after Mitotic Arrest via Phosphoregulation of Antiapoptotic Bcl-2 Proteins. Journal of Biological Chemistry 287, 39193–39204. 10.1074/jbc.M112.391854.

35. Lende-Dorn, B.A., Atkinson, J.C., Bae, Y., and Galloway, K.E. (2025). Chemogenetic tuning reveals optimal MAPK signaling for cell-fate programming. Cell Rep. 44, 116226. 10.1016/j.celrep.2025.116226.

36. Peterson, A.F., Ingram, K., Huang, E.J., Parksong, J., McKenney, C., Bever, G.S., and Regot, S. (2022). Systematic analysis of the MAPK signaling network reveals MAP3K-driven control of cell fate. Cell Syst. 13, 885–894.e4. 10.1016/j.cels.2022.10.003.

37. Gao, L., Fang, Y.-Q., Zhang, T.-Y., Ge, B., Tang, R.-J., Huang, J.-F., Jiang, L.-M., and Tan, N. (2015). Acidic extracellular microenvironment promotes the invasion and cathepsin B secretion of PC-3 cells. Int. J. Clin. Exp. Med. 8, 7367–7373.

38. Christensen, J., and Shastri, V.P. (2015). Matrix-metalloproteinase-9 is cleaved and activated by Cathepsin K. BMC Res. Notes 8, 322. 10.1186/s13104-015-1284-8.

39. Vidak, E., Javoršek, U., Vizovišek, M., and Turk, B. (2019). Cysteine Cathepsins and Their Extracellular Roles: Shaping the Microenvironment. Cells 8, 264. 10.3390/cells8030264.

40. Chung, C.Y.-S., Shin, H.R., Berdan, C.A., Ford, B., Ward, C.C., Olzmann, J.A., Zoncu, R., and Nomura, D.K. (2019). Covalent targeting of the vacuolar H+-ATPase activates autophagy via mTORC1 inhibition. Nat. Chem. Biol. 15, 776–785. 10.1038/s41589-019-0308-4.

41. Zhou, Y., Yang, X., Xu, W., Shen, S., Fan, W., Meng, G., Cheng, Y., Lu, Y., and Wei, Y. (2025). Rag GTPases control lysosomal acidification by regulating v-ATPase assembly in Drosophila. Journal of Biological Chemistry 301, 110400. 10.1016/j.jbc.2025.110400.

42. Ratto, E., Chowdhury, S.R., Siefert, N.S., Schneider, M., Wittmann, M., Helm, D., and Palm, W. (2022). Direct control of lysosomal catabolic activity by mTORC1 through regulation of V-ATPase assembly. Nat. Commun. 13, 4848. 10.1038/s41467-022-32515-6.

43. Wang, B., Martini-Stoica, H., Qi, C., Lu, T.-C., Wang, S., Xiong, W., Qi, Y., Xu, Y., Sardiello, M., Li, H., et al. (2024). TFEB–vacuolar ATPase signaling regulates lysosomal function and microglial activation in tauopathy. Nat. Neurosci. 27, 48–62. 10.1038/s41593-023-01494-2.

44. Iavazzo, M., Cinque, L., Levantovsky, S., Morrone, C., Monfregola, J., Raimondi, A., Polishchuk, E., De Cegli, R., Carrella, D., Nusco, E., et al. (2026). TFEB coordinates autophagosome biogenesis and ribophagy during starvation via SQSTM1. Sci. Adv. 12. 10.1126/sciadv.aea9302.

45. Franco-Juárez, B., Coronel-Cruz, C., Hernández-Ochoa, B., Gómez-Manzo, S., Cárdenas-Rodríguez, N., Arreguin-Espinosa, R., Bandala, C., Canseco-Ávila, L.M., and Ortega-Cuellar, D. (2022). TFEB; Beyond Its Role as an Autophagy and Lysosomes Regulator. Cells 11, 3153. 10.3390/cells11193153.

46. Fan, Y., Lu, H., Liang, W., Garcia-Barrio, M.T., Guo, Y., Zhang, J., Zhu, T., Hao, Y., Zhang, J., and Chen, Y.E. (2018). Endothelial TFEB (Transcription Factor EB) Positively Regulates Postischemic Angiogenesis. Circ. Res. 122, 945–957. 10.1161/CIRCRESAHA.118.312672.

47. Shin, S., Dimitri, C.A., Yoon, S.-O., Dowdle, W., and Blenis, J. (2010). ERK2 but not ERK1 induces epithelial-to-mesenchymal transformation via DEF motif-dependent signaling events. Mol. Cell 38, 114–127. 10.1016/j.molcel.2010.02.020.

48. Wang, K., Ji, W., Yu, Y., Li, Z., Niu, X., Xia, W., and Lu, S. (2018). FGFR1-ERK1/2-SOX2 axis promotes cell proliferation, epithelial–mesenchymal transition, and metastasis in FGFR1-amplified lung cancer. Oncogene 37, 5340–5354. 10.1038/s41388-018-0311-3.

49. Wojciechowski, M.C., Mahmutovic, L., Shu, D.Y., and Lovicu, F.J. (2017). ERK1/2 signaling is required for the initiation but not progression of TGFβ-induced lens epithelial to mesenchymal transition (EMT). Exp. Eye Res. 159, 98–113. 10.1016/j.exer.2017.03.012.

50. Astanina, E., Bussolino, F., and Doronzo, G. (2021). Multifaceted activities of transcription factor EB in cancer onset and progression. Mol. Oncol. 15, 327–346. 10.1002/1878-0261.12867.

51. Puertollano, R., Ferguson, S.M., Brugarolas, J., and Ballabio, A. (2018). The complex relationship between TFEB transcription factor phosphorylation and subcellular localization. EMBO J. 37. 10.15252/embj.201798804.

52. Roczniak-Ferguson, A., Petit, C.S., Froehlich, F., Qian, S., Ky, J., Angarola, B., Walther, T.C., and Ferguson, S.M. (2012). The Transcription Factor TFEB Links mTORC1 Signaling to Transcriptional Control of Lysosome Homeostasis. Sci. Signal. 5. 10.1126/scisignal.2002790.

53. Nowosad, A., and Besson, A. (2022). Lysosomes at the Crossroads of Cell Metabolism, Cell Cycle, and Stemness. Int. J. Mol. Sci. 23, 2290. 10.3390/ijms23042290.

54. Doronzo, G., Astanina, E., and Bussolino, F. (2021). The Oncogene Transcription Factor EB Regulates Vascular Functions. Front. Physiol. 12. 10.3389/fphys.2021.640061.

55. Astanina, E., Bussolino, F., and Doronzo, G. (2021). Multifaceted activities of transcription factor EB in cancer onset and progression. Mol. Oncol. 15, 327–346. 10.1002/1878-0261.12867.

56. Remy, D., Antoine-Bally, S., de Tocqueville, S., Jolly, C., Macé, A.-S., Champenois, G., Nemati, F., Brito, I., Raynal, V., Priya, A., et al. (2025). TFEB triggers a matrix degradation and invasion program in triple-negative breast cancer cells upon mTORC1 repression. Dev. Cell 60, 1018–1035.e8. 10.1016/j.devcel.2024.12.005.

57. Lee, D.H., Kim, T.M., Kim, J.K., and Park, C. (2019). ETV2/ER71 Transcription Factor as a Therapeutic Vehicle for Cardiovascular Disease. Theranostics 9, 5694–5705. 10.7150/thno.35300.

58. Mongiat, M., Andreuzzi, E., Tarticchio, G., and Paulitti, A. (2016). Extracellular Matrix, a Hard Player in Angiogenesis. Int. J. Mol. Sci. 17, 1822. 10.3390/ijms17111822.

59. Zeng, Z.-Z., Yao, H., Staszewski, E.D., Rockwood, K.F., Markwart, S.M., Fay, K.S., Spalding, A.C., and Livant, D.L. (2009). α5β1 Integrin Ligand PHSRN Induces Invasion and α5 mRNA in Endothelial Cells to Stimulate Angiogenesis. Transl. Oncol. 2, 8–20. 10.1593/tlo.08187.

