## Supplementary material for "V-ATPase-Driven Lysosomal Activation Orchestrates MEK2-Induced Endothelial Reprogramming": Figures S1, S2, S3 and Tables S1 and S2

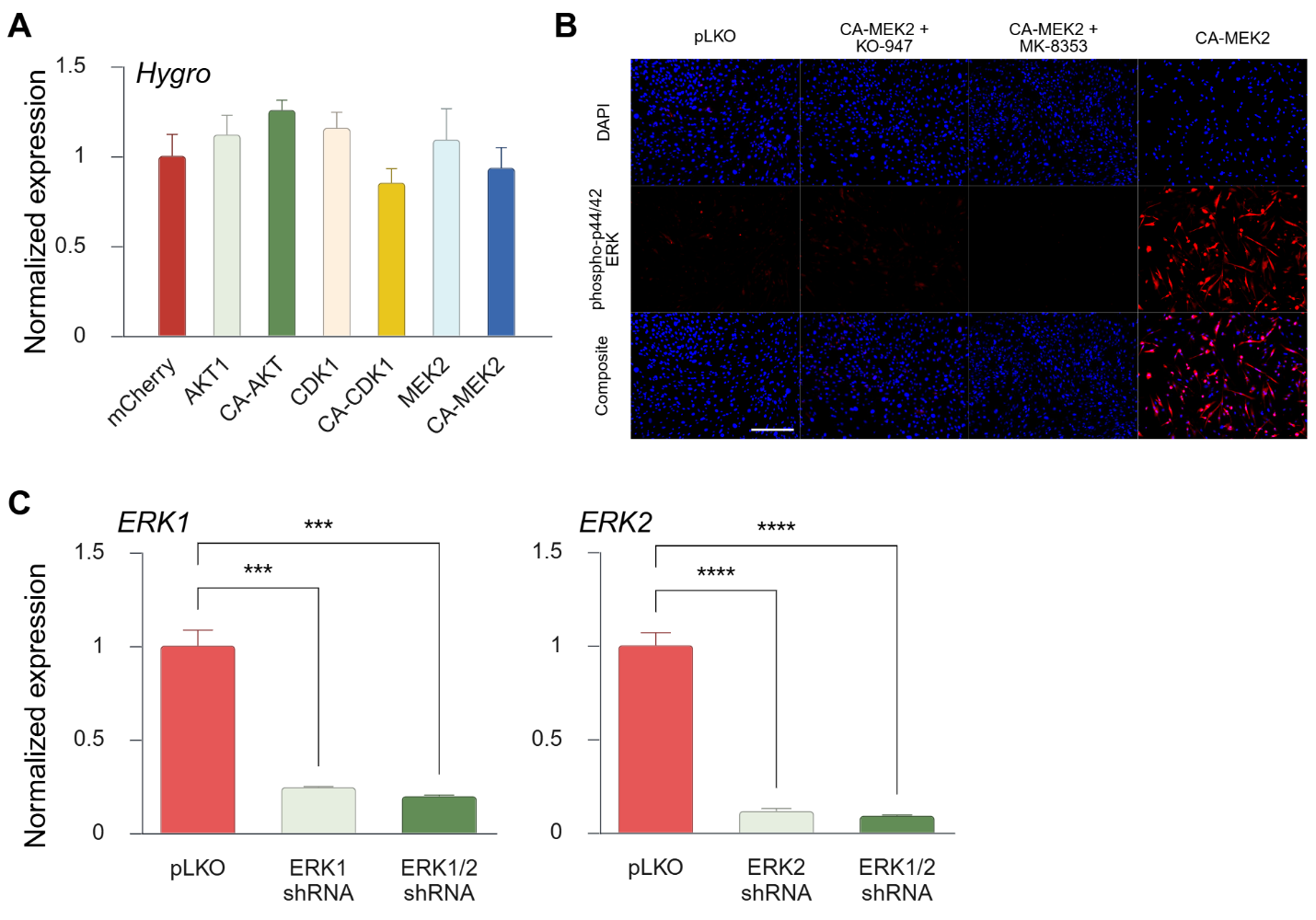

**Figure S1: Validation of ERK1/2 Activity and Functional Requirement for CA-MEK2–Induced Endothelial Reprogramming**

(A) Quantitative RT-PCR analysis showing relative expression of hygromycin resistance in MEFs transduced with wild-type or CA kinases AKT1, CDK1, and MEK2. Data represent three independent experiments, each with 3 technical replicates, and are shown as mean ± SEM.

(B) Immunofluorescence images of MEFs transduced with pLKO or CA-MEK2 for 9 days and treated with 10 μM KO-947 or MK-8353 for 2 days. Phospho-p44/42 ERK staining is shown in red; nuclei are counterstained with DAPI (blue). Scale bar, 100 μm.

(C) Quantitative RT-PCR analysis showing relative expression of *ERK1* and *ERK2* in response to MEFs transduced with *ERK1/2* shRNAs. ****p* < 0.001, and *****p* < 0.0001 vs. pLKO treatment transduction. Data represent three independent experiments, each with 3 technical replicates, and are shown as mean ± SEM.

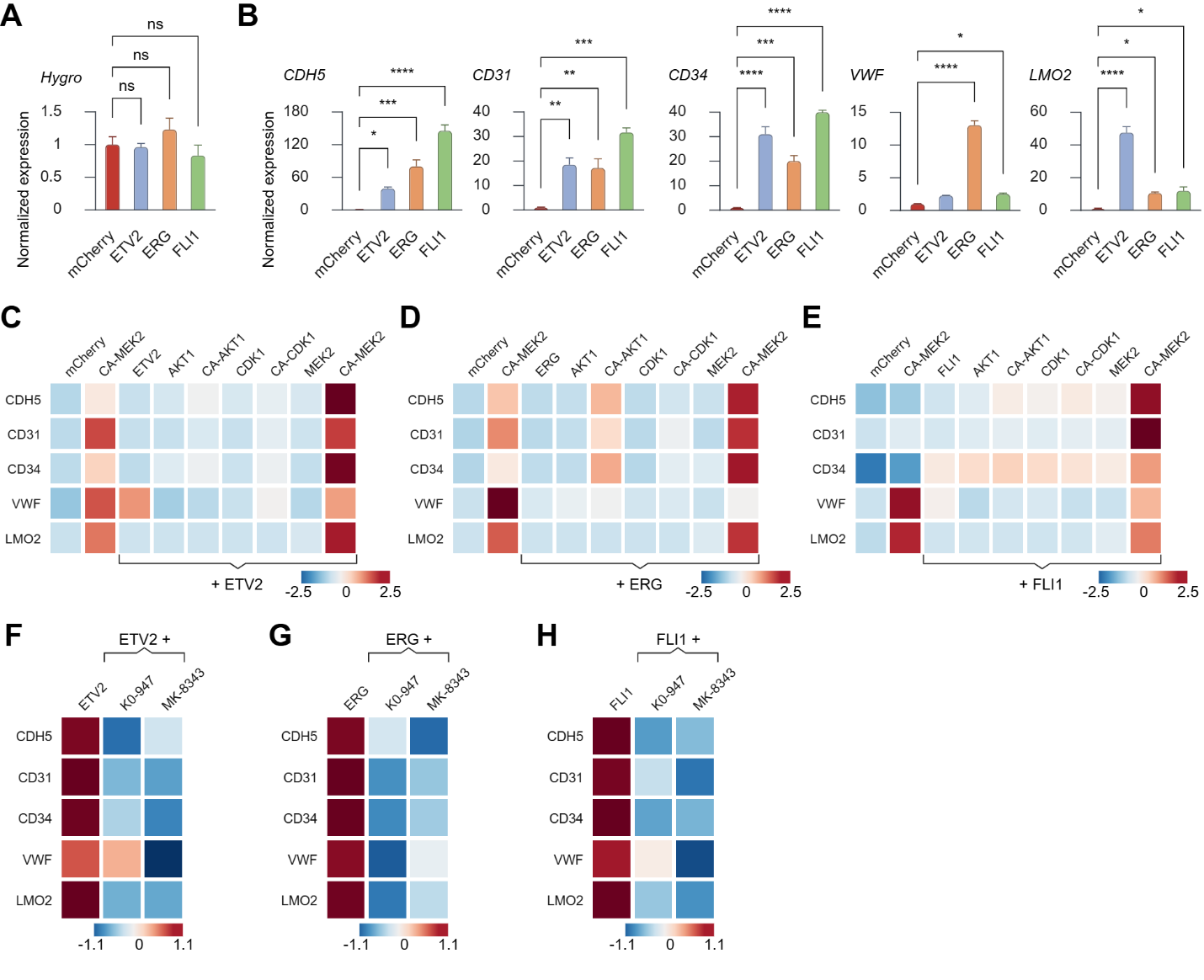

**Figure S2: CA-MEK2 Demonstrates the Strongest Capacity to Drive Endothelial Reprogramming among**

**the Candidate Kinases and Transcription Factors Tested.**

(A) Quantitative RT-PCR analysis showing relative expression of hygromycin resistance in MEFs transduced with ETV2, ERG, and FLI1. N = 3.

(B) Quantitative RT-PCR analysis showing relative expression of endothelial lineage genes *(CDH5, CD31, CD34, VWF,* and *LMO2)* in MEFs transduced with ETV2, ERG, and FLI1. Data represent three independent experiments and are presented as mean ± SEM. ∗*p* < 0.05, ***p* < 0.01, ****p* < 0.001, and *****p* < 0.0001 vs. mCherry transduction. Two-way ANOVA.

(C–E) Heatmaps of quantitative RT-PCR analysis showing relative expression of endothelial lineage genes *(CDH5, CD31, CD34, VWF,* and *LMO2)* in MEFs transduced with (C) ETV2, (D) ERG, or (E) FLI1, with or without co-transduction with CA-MEK2. N=3.

(F–H) Heatmaps of quantitative RT-PCR analysis showing relative expression of endothelial lineage genes *(CDH5, CD31, CD34, VWF,* and *LMO2)* in MEFs transduced with (F) ETV2, (G) ERG, or (H) FLI1, followed by treatment with 10 μM KO-947 or MK-8353 for 2 days. N=3.

Data represent three independent experiments of at least three technical replicates. Heatmap colors indicate relative expression levels, with red representing higher expression and blue representing lower expression.

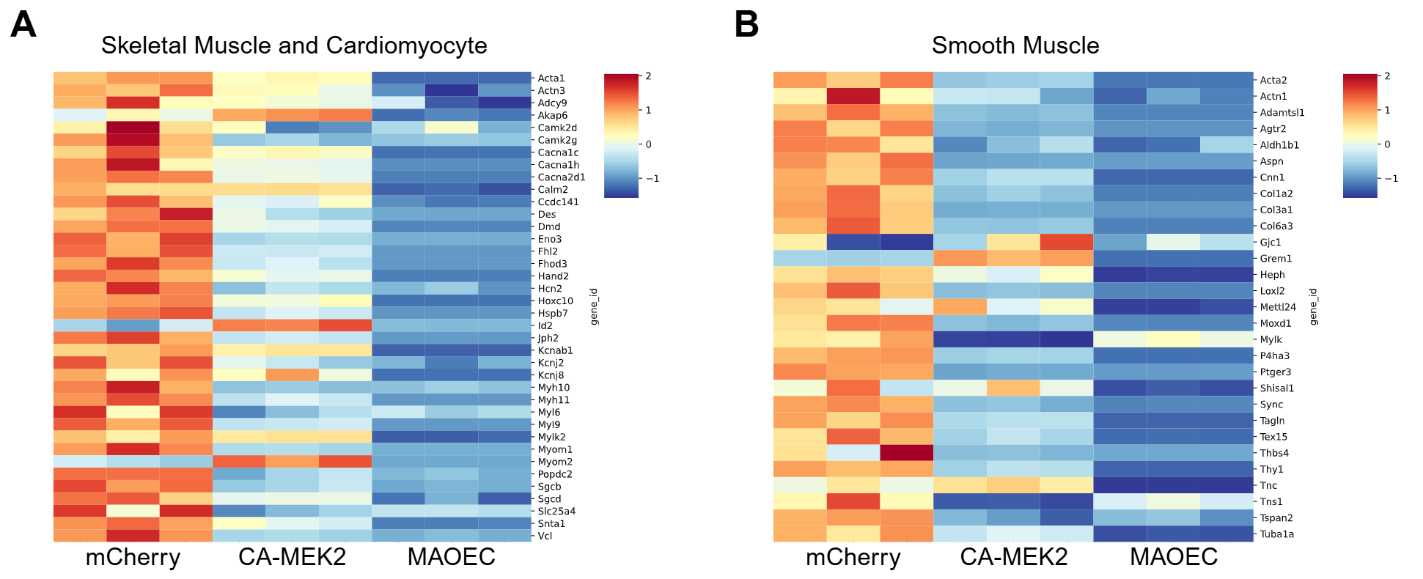

**Figure S3.** **CA-MEK2 transduction does not broadly activate skeletal, cardiac, or smooth muscle gene programs.**

(A and B) Heatmaps displaying RNA-seq–based expression of representative (A) skeletal muscle and cardiomyocyte genes and (B) smooth muscle–related genes in CA-MEK2–transduced MEFs. Gene expression was compared to mCherry-transduced controls. Data represent three biological replicates (N = 3).

**Table S1. The list of skeletal muscle and cardiomyocyte related genes selected for RNA sequencing analysis.**

| Acta1 | Actn3 | Adcy9 | Akap6 | Calm2 | Camk2d |
| --- | --- | --- | --- | --- | --- |
| Camk2g | Cacna1c | Cacna1h | Cacna2d1 | Ccdc141 | Des |
| Dmd | Eno3 | Fhl2 | Fhod3 | Hand2 | Hcn2 |
| Hoxc10 | Hspb7 | Id2 | Jph2 | Kcnab1 | Kcnj2 |
| Kcnj8 | Myh10 | Myh11 | Myl6 | Myl9 | Mylk2 |
| Myom1 | Myom2 | Popdc2 | Sgcb | Sgcd | Slc25a4 |
| Snta1 | Vcl |  |  |  |  |

**Table S2. The list of smooth muscle and related genes selected for RNA sequencing analysis.**

| Acta2 | Actn1 | Adamtsl1 | Agtr2 | Aldh1b1 | Aspn |
| --- | --- | --- | --- | --- | --- |
| Cnn1 | Col1a2 | Col3a1 | Col6a3 | Gjc1 | Grem1 |
| Heph | Loxl2 | Mettl24 | Moxd1 | Mylk | P4ha3 |
| Ptger3 | Shisal1 | Sync | Tagln | Tex15 | Thbs4 |
| Thy1 | Tnc | Tns1 | Tspan2 | Tuba1a |  |
